# Proton Coupling and the Multiscale Kinetic Mechanism of a Peptide Transporter

**DOI:** 10.1101/2021.09.10.459748

**Authors:** Chenghan Li, Zhi Yue, Simon Newstead, Gregory A. Voth

## Abstract

Proton coupled peptide transporters (POTs) are crucial for the uptake of di- and tri-peptides as well as drug and pro-drug molecules in prokaryotes and eukaryotic cells. We illustrate from multiscale modeling how transmembrane proton flux couples within a POT protein to drive essential steps of the full functional cycle: 1) protonation of a glutamate on transmembrane helix (TM) 7 opens the extracellular gate, allowing ligand entry; 2) inward proton flow induces the cytosolic release of ligand by varying the protonation state of a second conserved glutamate on TM10; 3) proton movement between TM7 and TM10 is thermodynamically driven and kinetically permissible via water proton shuttling without the participation of ligand. Our results, for the first time, give direct computational confirmation for the alternating access model of POTs, and point to a quantitative multiscale kinetic picture of the functioning protein mechanism.

**SIGNIFICANCE:** Proton-coupled peptide transporters (POTs) utilize transmembrane proton gradient to deliver small peptides and peptide-like drug molecules into cells. Despite extensive biochemical and structural studies, major question regarding protonation-induced shift from inward-facing state to outward-facing state remains obscure. Here, we report direct evidence through multiscale simulations that the extracellular salt bridge controls the outward-open conformational transition of POTs, and how proton influx through POTs couples ligand transport. The computational modeling also suggests a multiscale kinetic mechanism of POTs.

## INTRODUCTION

Proton coupled oligopeptide transporters (POTs) utilize a membrane pH gradient to drive cellular uptake of di-/tri-peptides and their analogs, with homologs found in all bacterial and eukaryotic genomes (1,2). Mammalian cells contain two POT proteins, PepT1 (SLC15A1) and PepT2 (SLC15A2), which are responsible for the bulk uptake and retention of dietary peptides in the small intestine and kidneys, respectively (1). PepT1 and PepT2 can also transport prodrug molecules and are increasingly recognized as important targets for rational drug design to improve drug pharmacokinetics (3-5). The POT proteins use an alternating access mechanism, where peptide transport is realized through conformational switching between two major conformations, termed inward-facing (IF) and outward-facing (OF) states (6). An occluded (OC) state has also been observed (7,8), although this is likely to be transitory. In OF and IF conformations, the ligand-binding site is accessible from the extracellular or the intracellular environment, while access is prohibited from both sides in the OC state (9). The alternating access cycle in POTs has been determined following structural and biochemical studies on bacterial homologs (10) and is rationalized on the basis of alternating formation and breaking of conserved salt bridge interactions, which drive the structural changes following peptide and proton binding (10,11).

Several conserved side chains have been identified that play key roles in the transport mechanism. In particular, it was found that a salt bridge between a highly-conserved glutamate/aspartate on transmembrane helix (TM) 7 and the arginine/lysine on TM1 stabilizes the closed state of the extracellular gate (**Fig. 1**) (6,12,13). Protonation of the glutamate/aspartate was predicted to break the salt bridge and trigger the IF-to-OF conformational change, allowing ligand access from the extracellular side (6,11), but direct evidence for this crucial part of the transport mechanism is still lacking. Molecular dynamics (MD) simulations have been used to investigate the protein and ligand dynamics and coupling in POTs (11,14-18). However, many MD-based studies contain contradictory findings, ascribing different functions for these side chains in different POT family homologs and highlighting the challenges of sufficient sampling of the extended phase space spanned by protein, ligand, protons, and their associated hydration. It was reported in an MD study on GkPOT from *Geobacillus kaustophilus*, for example, that the charge state of the TM7 Glu does not affect the conformational state of the transporter (15). A later study that employed enhanced conformational sampling found the OF conformation of PepT_So_ from *Shewanella oneidensis* to be more stable than the IF one when the TM7 Glu is deprotonated (16), suggesting the TM1-TM7 salt bridge cannot determine the extracellular gate closure. A more recent computational study systematically evaluated the impact of the TM7 Glu and ligand on the conformational free energy landscape of PepT_St_ from *Streptococcus thermophilus*, and found the IF and OC states were the most stable, independent of the TM7 Glu and ligand (17), inconsistent with the expectation for a stable apo OF state to allow efficient access to ligands in the periplasmic space. These studies were conducted in a fixed charge and non-reactive (fixed chemical bonding topology) manner that limited their ability to fully sample the complete transport cycle and especially the proton coupling mechanism.

**FIGURE 1.**
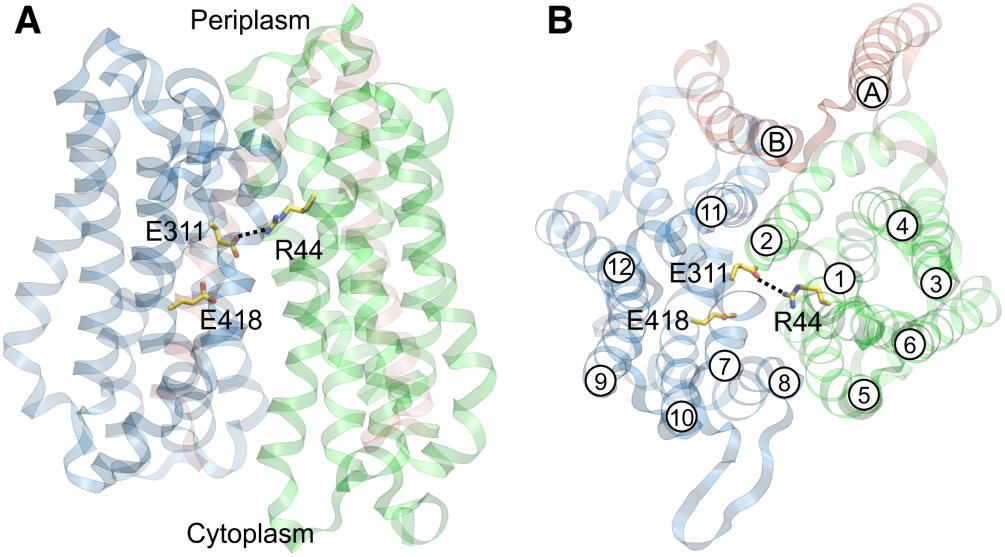
PepT_Sh_ structure (PDB: 6EXS) and essential ionizable residues. **(A)** Side view parallel to the cellular membrane. The N-terminal bundle is rendered in green, the C-terminal bundle is in blue while flanking helices A and B are in red. Black dashed line indicate a salt bridge. **(B)** Top view from periplasm, perpendicular to the cellular membrane. The helices are indicated with labels in cycles.

Previously, we discovered a homolog of PepT1 from the bacterium *Staphylococcus hominis*, termed as PepT_Sh_, transports a peptide-like thioalchohol precursor, cysteinylglycine-3-methyl-3-sulfanylhexan-1-ol (S-Cys-Gly-3M3SH) (19). The compound is secreted by the apocrine gland, mostly in the armpit and groin regions of the body (20). Uptake of the S-Cys-Gly-3M3SH ligand into *S. hominis* results in its biotransformation into an odorous volatile responsible for human body odor (19). PepT_Sh_ shares high sequence similarity with PepT1 and PepT2 and can transport prodrugs such as valacyclovir and 5-aminolevulinic acid (19,21,22). Hence, PepT_Sh_ represents an ideal model for understanding the proton coupling mechanism across the bacterial and human homologs within the POT family. Our previous work captured the IF state of PepT_Sh_ bound with S-Cys-Gly-3M3SH as well as several drugs (19,21), although the OF and OC states were not captured. It therefore remains challenging to complete the functional cycle of proton-coupled ligand transport starting from static IF structures.

In this work, we performed extensive all-atom MD and multiscale reactive MD simulations (MS-RMD; previously termed multistate empirical valence bond or MS-EVB) (23,24) combined with enhanced free energy sampling to elucidate how proton transport (PT) through key residues drives conformational changes and ligand translocation. We have demonstrated here that it is indeed the titrations on the TM7 and TM10 glutamates that trigger the global conformational change and ligand release in PepT_Sh_. Extending previous studies, we have also evaluated the energetics associated with proton movements and found the PT free energies are subtly modulated by the transporter conformational state and the bound ligand. Our findings, which are consistent with a previous study looking at extracellular gate dynamics in PepT_Xc_, a POT family homolog from *Xanthomonas campestris* (11), reveal a coupled and cooperative multiscale kinetic mechanism between the transporter conformation, proton, and ligand motions, which represents a major step toward a complete and quantitative description of the full transport cycle.

## METHODS

### Classical molecular dynamics simulations

The MD computational model was constructed using the CHARMM-GUI (25,26) from the holo IF crystal structure of PepT_Sh_ (PDB: 6EXS) by embedding the protein into a 90-Å × 90-Å bilayer composed of 1-palmitoyl-2-oleoyl-*sn*-glycero-3-phosphocholine (POPC) lipids and then solvating the system with a 20-Å layer of TIP3P water (27) on both sides of the membrane with 0.15 M sodium chloride. The protonation states of ionizable residues were assigned according to p*K*_a_ predictions from constant-pH MD (CpHMD) simulations (detailed in below) and are listed in **Table S1**. The convergence of CpHMD p*K*_a_ calculations for crucial residues are shown in **Fig. S9**. The CHARMM 22/CMAP (28-30) force field was employed to describe the protein and the CHARMM36 (31) was used for lipid interactions. The ligand was modeled as a dipeptide consisting of S-Cys-3M3SH (CSM) and a glycine. CSM was added as a non-standard amino acid to the force field, and was parameterized by CHARMM General Force Field (CGenFF, version 4.0) (32) using ParamChem (version 2.2.0). The CGenFF parameters for the backbone and sidechain atoms up to S^*γ*^ were replaced by those for cysteine from CHARMM22 force field. The atomic charge of *C*^*β*^ was adjusted for charge neutrality (Appendix I). Particle mesh Ewald (PME) (33,34) with a cutoff of 12.0 Å and a precision of 10^−5^ was used to compute the electrostatic interactions. The Lennard-Jones (LJ) non-bonded interactions was force-switched from 10 Å to 12 Å. The system was equilibrated following the CHARMM-GUI protocol (35), followed by a 200-ns additional equilibration with 1000 kJ/nm^2^ harmonic restraints on protein heavy atoms. In the additional equilibration and production runs, the system was integrated by the leap-frog algorithm with a 2-fs long time step, and all the bonds involving hydrogen atoms were restrained using the LINCS algorithm (36). The temperature was controlled at 303.15 K by the Nosé-Hoover thermostat (37,38) and the pressure was controlled at 1 atm by the Parrinello-Rahman semiisotropic barostat (39). The simulation time and the initial configurations of production runs were summarized in **Table S1**. All classical MD simulations were conducted in the GROMACS package (40).

### Multiscale reactive molecular dynamics (MS-RMD) simulations

The MS-RMD approach was well documented in refs (23,24). In brief, the MS-RMD approach provides an efficient way to simulate molecular systems with explicit modeling of chemical reactions (varying bonding topology in time). This was achieved by considering the system as a linear combination of diabatic states {|*i*⟩}, each of which corresponds to a different bonding topology. The Hamiltonian of the system is then expressed in the following diabatic state representation:

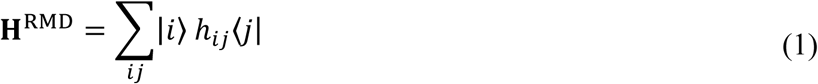

The diagonal term *h*_*ii*_ is taken to be the energy function described by the classical force field, namely the CHARMM22/CMAP for proteins and CHARMM36 for lipids. The off-diagonal element, *h*_*ij*_, is modeled by a physically inspired ansatz in a MM form. The detailed definitions of these terms are provided in the ref (41). The ground state of the reactive system can be obtained through solving the following secular equation “on the fly” as a function of nuclear configuration, such that

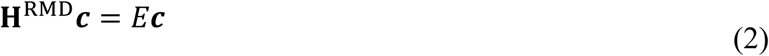

The eigenvector ***c*** = {*c*_*i*_} with the lowest eigen energy is the adiabatic wave-function of the ground state. The atomic forces, as the energy gradient, are *c*_*i*_ weighted diabatic forces according to the Hellmann-Feynman theorem:

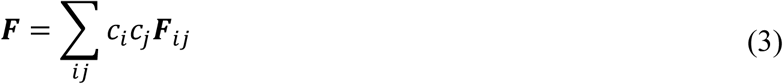

The diabatic matching approach (41) was used to parametrize the ionizable MS-RMD model for glutamates described by the CHARMM22 force field and the parameters are summarized in **Table S2**.

The MS-RMD simulations were initiated from classical MD equilibrations. The electrostatics was computed by the particle-particle particle-mesh method (42) with a cut-off of 10 Å and an accuracy criterion of 10^−4^. The non-bonded LJ potential was energy-switched from 8 Å to 10 Å. The system was integrated by the Nosé–Hoover chain thermostat to maintain a 303.15 K temperature in the NVT ensemble using a time step of 1 fs. The MS-RMD simulations were performed by the LAMMPS MD package (43) coupled to RAPTOR (44) to enable chemical reactions.

### Enhanced free energy sampling and rate calculations

The PT between E311 and E418 in the apo form was enhanced by the well-tempered metadynamics (WT-MTD) approach (45). The CV used in the WT-MTD was defined as a distance ratio:

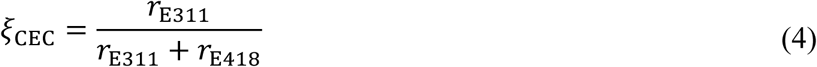

where *r*_E311_ and *r*_E418_ are the minimum distance between the center of excess charge (CEC) and carboxyl oxygen atoms of E311 or E418. Due to the charge delocalized nature of a hydrated excess proton, its effective position is tracked by the center of excess charge (CEC) introduced by the excess proton defect, or equivalently, the “electron hole” created by the extra proton nucleus. In MS-RMD, the CEC is defined as the *c*_*i*_ weighted center of charge (COC) of the proton-carrier species in each diabatic state (46),

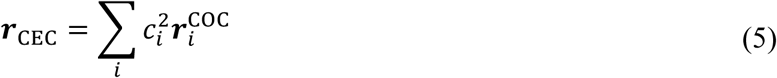

The minimum distance between CEC and carboxyl oxygens was implemented by a softmin function, defined as

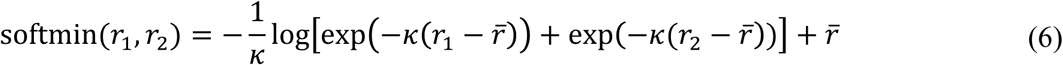

where *κ* = 40 Å^−1^, *r*_1_ and *r*_2_ denote the CEC separation from the two carboxyl oxygen atoms, and 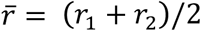. The initial Gaussian height in WT-MTD was 0.8 kcal/mol and scaled according to a bias factor of 12. The Gaussians were deposited on the *ξ*_CEC_ dimension every 1 ps with a fixed width of 0.01. In both apo IF and apo OF states, two replicates of metadynamics were run for at least 10 ns.

The PT between E311 and E418 in holo IF state was sampled by umbrella sampling (47) owing to its convenience in distributing sampling tasks on multiple computer nodes. The umbrella window centers were placed from *ξ*_CEC_ = 0 to *ξ*_CEC_ = 1 every 0.025 with harmonic force constants ranging from 2000 kcal/mol to 3500 kcal/mol. The simulation time for each window ranged from 370 ps to 7 ns depending on its convergence, resulting in a 57-ns simulation time in total. The enhanced sampling functionality was provided by PLUMED 2 (48) coupled to LAMMPS MD engine and the RAPTOR MS-RMD software.

The free energy surfaces (potentials of mean force; PMFs) were computed from metadynamics or umbrella sampling data using the dynamic histogram analysis method (DHAM) (49), and so was the Markov transition matrix in the CV space. The CV-position-dependent diffusion constants *D*(*ξ*) were computed following the same procedure in Refs. (50,51) using the transition matrix, and the reaction rate constant was computed as the inverse of the mean first passage time (MFPT) (52),

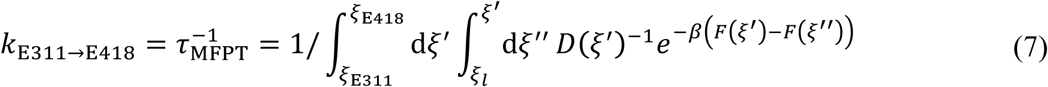

assuming Smoluchowski dynamics of for the variable *ξ*_CEC_. In eq 7, the *ξ*_E311_ and *ξ*_E418_ are the CV values that correspond to free energy minima of protonated E311 and E418 respectively, *ξ*_*l*_ is the CV value that corresponds to the lower boundary of the E311 free energy well, *β* = 1/*k*_*B*_*T* is the inverse temperature, and *F* (*ξ*) is the PMF. The reverse PT rate was computed via the detailed balance relation

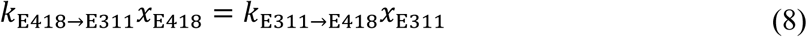

where *x* is equilibrium mole fraction computed as the integral of Boltzmann factor *e*^−*βF*(*ξ*)^ in the E311 basin or the E418 basin. The errors reported for WT-MTD PMFs and PT rates were computed as the standard deviation between two replicates, while the errors in umbrella sampling counterparts were computed from the standard deviation of the last 5 blocks of equally partitioning the trajectories into 6 blocks.

### Characterization of gate sizes and gate hydration

The extracellular and intracellular gate sizes were calculated as the tip distance between TM1,2 and TM7,8 and between TM4,5 and TM10,11. We first defined four virtual atoms as backbone geometric centers of (1) residues 46–52 and 64–70, (2) residues 316–324 and 342–347, (3) residues 140–146 and 154–160, and (4) residues 426–432 and 438–444, to represent the tip positions of TM1,2, TM7,8, TM4,5 and TM10,11 respectively. The extracellular gate size was then defined as the distance between virtual atoms (1) and (2), and the intracellular gate size was similarly defined as the distance between virtual atoms (3) and (4).

The intracellular gate water density was defined as the number of water molecules in a quadrangular prism divided by its volume. The prism base was defined as the *xy*-plane (membrane plane) projection of the tip residues of TM4, 5, 10, and 11, represented by four virtual atoms defined as the backbone geometric center of residues 140–146, 154–160, 426–432 and 438–444. The height of the prism was defined as the *z* range of the backbone atoms used to define the virtual centers.

### CpHMD simulation

To enable titration, doubly protonated HIS residues were used, dummy hydrogen atoms were added to ASP, GLU, and C-terminus (CT) carboxylates in *syn* positions. The membrane-enabled (53) hybrid-solvent Constant pH MD (CpHMD) (54) simulations were conducted using CHARMM (55) (version c42b2) with the pH-based replica exchange (pH-REX) enhanced-sampling protocol (54). The simulations represented protein, lipids, and waters using the CHARMM22/CMA force field, the CHARMM36 model, the CHARMM-modified TIP3P model, respectively. The CpHMD parameters for the N-terminus (NT) (Appendix II and **Fig. S10**) and CT (Appendix III and **Fig. S11**) were derived using the protocols described by Lee *et al*. (56) and Khandogin *et al*. (57), respectively. The LJ interactions were force-switched (58) from 8 to 12 Å. Electrostatic interactions were computed using the PME method with a real-space cutoff of 12 Å and a sixth-order interpolation with 1-Å grid spacing. The non-bonded neighbor list was updated heuristically. To allow for a 2-fs step length, SHAKE (59) was used to restrain the bonds connecting hydrogen atoms. All simulations were conducted with periodic boundary conditions at 303.15 K by the Nosé-Hoover thermostat, and 1 atm by the Langevin piston pressure-coupling algorithm (60).

In CpHMD, a fictitious titration coordination λ that updates simultaneously with the spatial coordinates is coupled to each ionizable site to describe its charge state. The λ particles with a mass of 10 atomic mass unit are propagated using the extended Langevin algorithm (61) with a collision frequency of 5 ps^−1^ and updated every 10 MD steps to allow for water relaxation. The hybrid-solvent CpHMD uses the leapfrog Verlet integrator (62) to propagate the conformational dynamics in explicit solvent and lipid molecules. The implicit solvent and membrane modelled by the generalized-Born (GB) model GBSW (63,64) with optimized GB input radii (65) is used to calculate the electrostatic hydration forces on λ particles. An infinite low-dielectric-constant (*ε* = 2) slab with a high-dielectric-constant (*ε* = 80) exclusion cylinder aligned with the membrane normal is used to represent protein-embedded membrane in GBSW. The implicit-membrane had a thickness of 35 Å and the radius of the exclusion cylinder was 25 Å. The dielectric constant (*ε*) was switched from 2 to 80 within 2.5 Å from both membrane surfaces. An ionic strength of 0.150 M was used for the Debye–Hückel term (66) in GB. All Asp, Glu, His, Lys, Arg, Cys, Tyr residues, as well as the NT and CT of the ligand were allowed to ionize. A cylindrical restraint with a force constant of 1 kcal/(mol•Å^2^) was applied to the center of mass of the protein heavy atoms via the MMFP utility in CHARMM (55) to prevent the protein from lateral drift. Ions were excluded from the hydrophobic membrane region (–16.5 Å < Z < 16.5 Å) by a planar restraint of 5 kcal/(mol•Å^2^) via MMFP. In pH-REX, exchanges between neighboring pH replicas were attempted every 500 MD steps. Decision of accepting or refusing an exchange attempt was determined using the Metropolis criterion (67). Two independent runs were conducted for apo-PepT_Sh_: one with and the other without titrations on Lys/Arg/Cys/Tyr residues (**Table S3**). The 1^st^ apo run placed 40 replicas from pH 2.0 to 11.75 with an interval of 0.25 and lasted 20 ns per replica. The 2^nd^ apo run placed 40 replicas from pH 2.0 to 9.0 with an interval of 0.125 or 0.25 and lasted 20 ns per replica. The apo run of dipeptide CSM-GLY (i.e., ligand) placed 20 replicas from pH 7.0 to 16.5 with an interval of 0.5 and lasted for 5 ns. The holo run (i.e., PepT_Sh_ with CSM-GLY) placed 40 replicas from pH 2.0 to 11.75 with an interval of 0.25 and lasted 10 ns per replica.

### Calculation of p*K*_a_

The p*K*_a_ was computed by fitting the deprotonated fraction *S*^unprot^ *vs*. pH to the Hill equation (68),

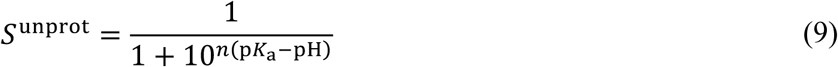

where the Hill coefficient *n* describes the steepness of the transition region in a titration curve. *S*^unprot^ was determined by counting the population of protonated (defined as those with *λ* ≤ 0.1) and deprotonated (*λ* ≥ 0.9) states for every pH.

## RESULTS AND DISCUSSION

### TM10 Glu is crucial for ligand binding and its titration controls ligand release

Our earlier PepT_Sh_ crystal structure (PDB: 6EXS) shows the transporter coupled to S-Cys-Gly-3M3SH in the IF state (**Fig. 2A**). The hydrophobic tail of the ligand was discovered to be inserted into a pocket created by TM7 and TM10 (**Fig. 2B**). Although a TM10 Glu (E418) is not directly coordinated to the ligand in the crystal structure, mutagenesis of the Glu in PepT_Sh_, PepT_St_ (6) and human PepT1 (69) has suggested the Glu is important for proton coupling and ligand recognition. This prompted us to run two sets of simulations from the crystal structure: E33^H^/E311^−^/E418^H^ (run #1) and E33^H^/E311^−^/E418^−^ (run #2). We defined a collective variable (CV) as the minimal distance between the TM9 backbone atoms and the ligand heavy atoms to assess the binding of the ligand tail in the pocket (see **Fig. 2B**). We found that if E418 is protonated, the simulations could reproduce the crystal ligand binding pose, which features the ligand tail packed in the TM7-TM10 pocket, as demonstrated by the time evolution and distribution of the distance CV (**Fig. 2C**, blue lines). When E418 is deprotonated, on the other hand, the ligand exits the TM7-TM10 binding pocket (**Fig. 2C**, red lines), yielding a lower binding posture that differs somewhat from the crystal (**Fig. 3A**). These results are consistent with the optimal conditions for the ligand bound crystal structure of pH 5.0. Replicated runs further indicate that the binding mode captured by the crystal structure is not rigid, since the ligand tail may also depart the pocket when E418 is protonated (**Fig. S1**, left panels, replica 3), while a deprotonated E418 can accelerate this process (**Fig. S1**, right panels).

**FIGURE 2.**
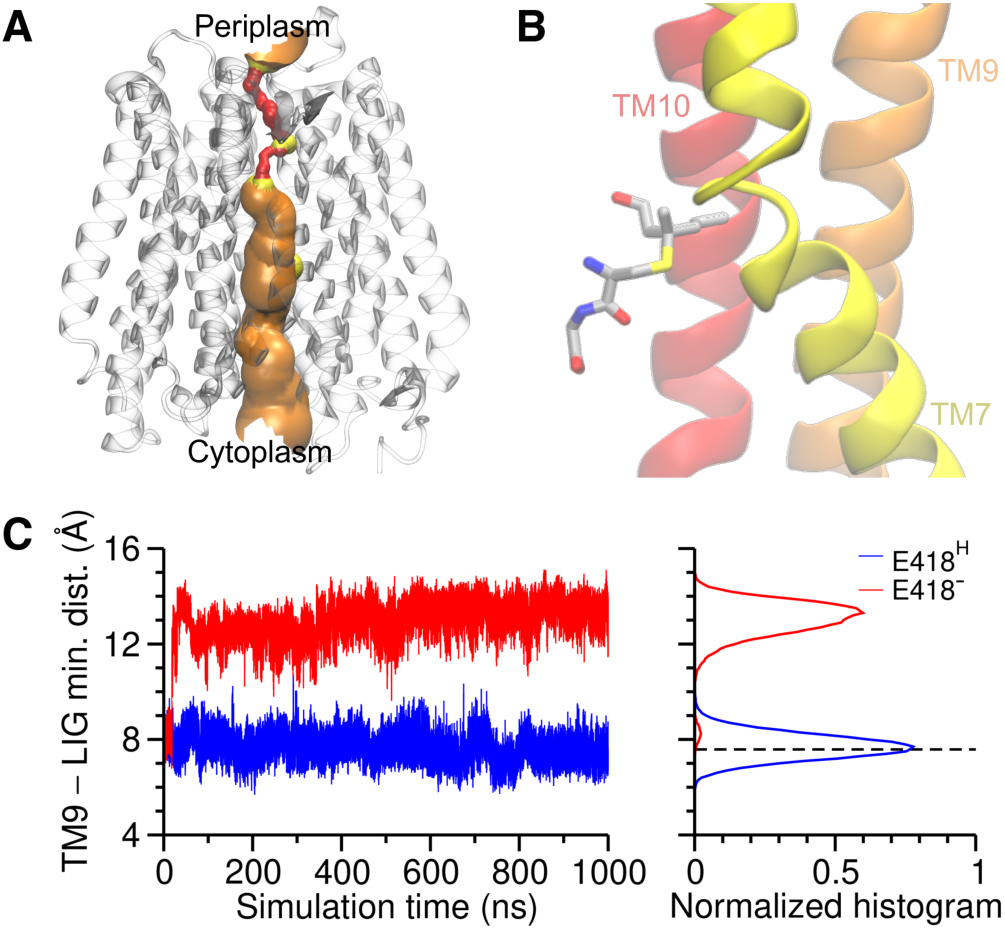
Role of E418 in ligand binding. **(A)** The inward-facing holo crystal structure (PDB: 6EXS). The pore radius profile was computed by the HOLE program (70). The region that forbids water (pore radius < 1.15 Å) is colored red, the region that allows single water permeation (1.15 Å < pore radius < 2.30 Å) is colored yellow, and orange indicates pore radius > 2.30 Å. This color scheme will be used for the following molecular figures. **(B)** The position of the ligand and the TM7-TM10 pocket as well as TM9 in the crystal structure. **(C)** The minimum distance between the ligand and TM9 backbone atoms in E33^H^/E311^−^/E418^H^ (run #1-1) and E33^H^/E311^−^/E418^−^ (run #2-1) simulations. The value in the crystal structure was indicated by the dashed horizontal line.

**FIGURE 3.**
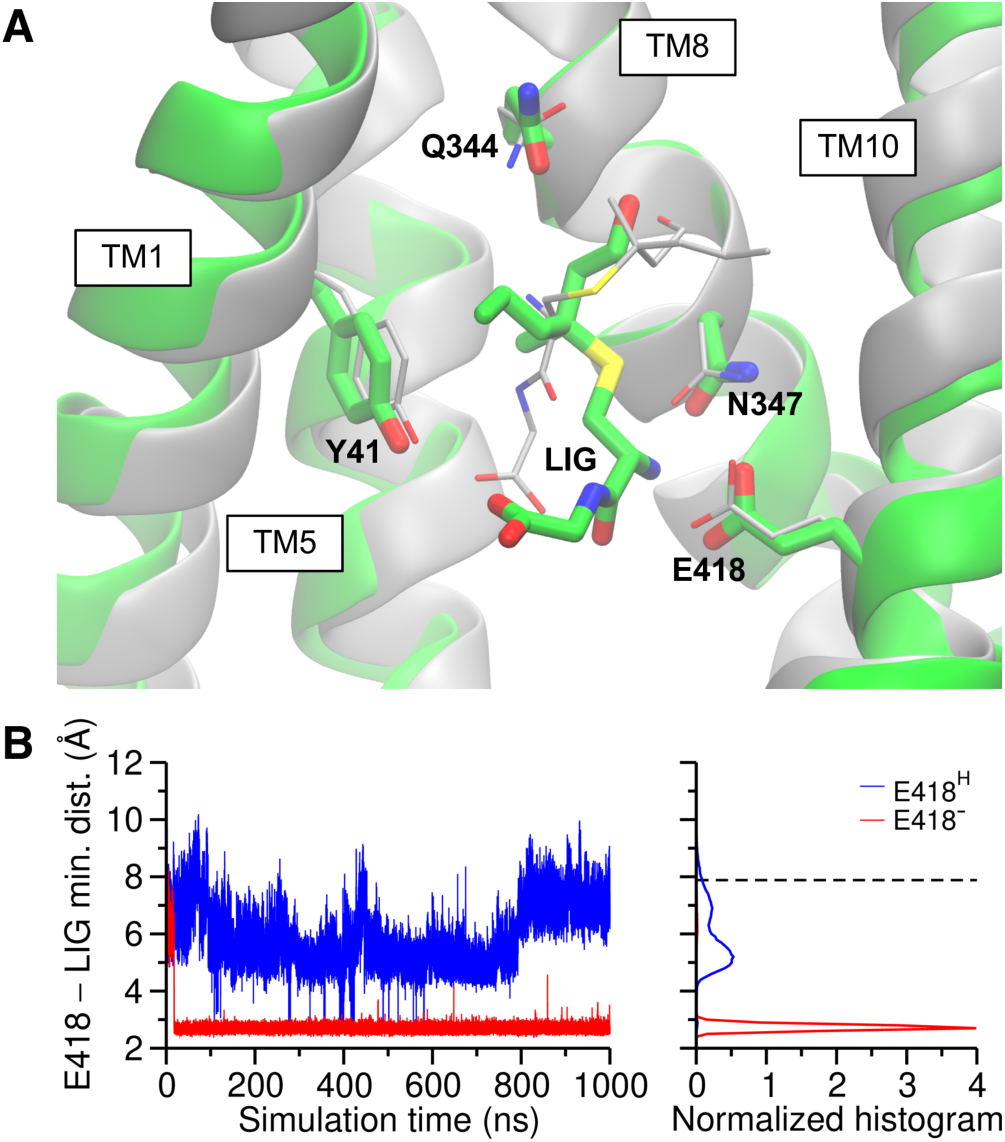
The binding mode of the ligand when E418 is deprotonated. **(A)** A super-position of the crystal structure (grey) and the equilibrated structure with a deprotonated E418. **(B)** The minimum distance between the E418 carboxyl and the ligand N-terminus in E33^H^/E311^−^/E418^H^ (run #1-1) and E33^H^/E311^−^/E418^−^ (run #2-1) simulations. The value in the crystal structure is indicated by the dashed horizontal line.

To further understand the role of E418 in ligand binding, we computed the minimal distance between the carboxyl of E418 and the N-terminus of the ligand. The protonated E418 simulations (run #1), which serve as a control, faithfully reproduce the weak ligand–E418 interaction seen in the crystal structure, as evidenced by the crystal distance value falling within the MD distribution measured (**Fig. 3B**, blue lines; **Fig. S2**, left panels). A deprotonated E418 (run #2), however, interacts with the ligand quickly (∼50 ns) by forming a salt bridge (**Fig. 3B**, red lines; **Fig. S2**, right panels), causing the ligand tail to leave the TM7-TM10 pocket as previously stated. Interestingly, despite these discrepancies in its binding pose, the bound ligand remains stable in the binding site over the course of the 3.5-*μs* MD simulation, possibly due to the stability gained from the salt bridge with E418 compensating for the loss from leaving the TM7-TM10 pocket. E418 is well-solvated in the IF conformational state, and the water network fully connects the Glu to the cytosolic bulk, so a pH gradient should be able to drive the proton of E418 down to the cytosol. As such, the crystal structure could represent a prelude to the deprotonated E418 state, in which the ligand transfers into the lower binding location after the proton dissociates from E418.

We next modeled the PT driven by the pH gradient from the periplasm to the TM10 Glu, and initiated E33^H^/E311^−^/E418^H^ (run #3) simulations from the equilibrated configurations sampled in E33^H^/E311^−^ /E418^−^ (run #2) simulations. We will validate the feasibility of this hypothetical PT in following sections, but we now report the consequences of this protonation state change. As shown in **Fig. 4A**, protonation of E418 makes the ligand unstable and eventually triggers the ligand release into the cytosol. Notably, the salt bridge between E418 and the ligand breaks almost instantly upon the protonation, but the ligand remains near the binding site for several hundreds of nanoseconds. During this metastable period, we looked at the interacting residues with the ligand (**Fig. 4C**) and compared them to the interaction pattern in the stable ligand-bound state when E418 is deprotonated (**Fig. 4B, Fig. S3**). We found that the protonation indeed weakens the interaction between the ligand and E418, and as a result, the Q344 becomes the dominant residue interacting with the ligand via a hydrogen bond to its sidechain hydroxyl group. The missing hydroxyl in di-/tri-peptides could be one of the reasons why S-Cys-Gly-3M3SH is transported at a slower rate than these other ligands (19). The lack of contacts with E418 and N347 (**Fig. 4D**) resulted in faster ligand release in a replicate than the other two **(Fig. 4A** red vs. green and blue), emphasizing the importance of the residues for ligand binding, which is consistent with the mutagenesis results that E418A and N347A decreased or eliminated transport efficiency (19).

**FIGURE 4.**
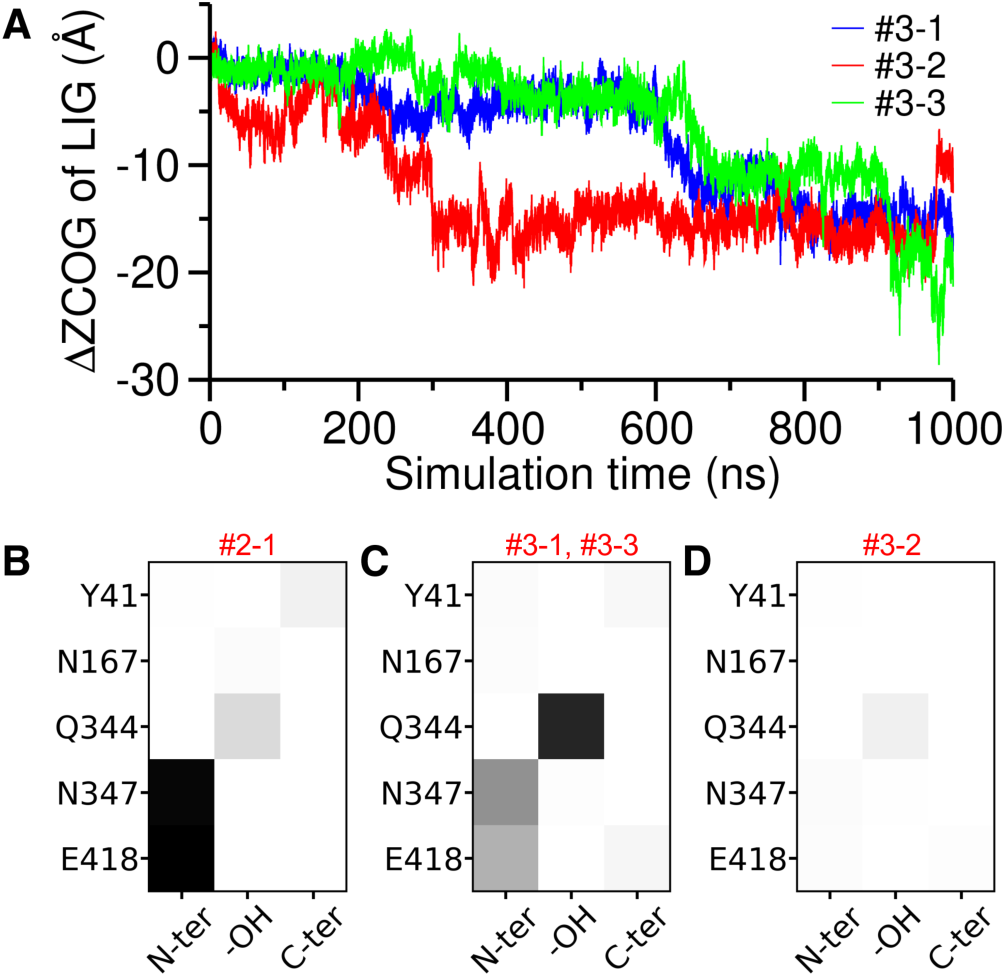
Proton-induced ligand release in E33^H^/E311^−^/E418^H^ (run #3) simulations. **(A)** Time evolution of the *z* coordinate of ligand geometric center with respect to the center of the membrane (ΔZCOG). **(B)** The contact map between ligand functional groups and binding-site residues (B) in the bound state with a deprotonated E418 (run #2-1), **(C)** in the first 200-ns metastable state in runs #3-1 and #3-3, and **(D)** in the first 20-ns of run #3-2.

### Titration of TM7 Glu triggers conformational change to outward-facing (OF) state

The highly conserved TM7 glutamate/aspartate–TM1 arginine/lysine pair and the TM7 serine–TM2 histidine pair are two of the critical salt bridge and hydrogen bond interactions that maintain the closure of the POT extracellular gate. We have previously shown (11) that the extracellular gate of PepT_Xc_ is regulated by the interaction between the TM2 His and TM7 Ser, and the protonation of His opens the gate by disrupting the His–Ser hydrogen bond. The His and Ser residues are conserved in PepT_Xc_ and PepT_So_, a POT from *Shewanella oneidensis*, as well as mammalian POTs; however, the TM2 His is absent from certain bacterial POTs, such as PepT_St_, and PepT_Sh_ studied in this work. Instead, the salt bridge between TM7 Glu/Asp and TM1 Arg/Lys, which can be seen in virtually every IF structure of POTs, might serve as an alternate mechanism for controlling the extracellular gate. We performed E33^H^/E311^−^/E418^−^ simulations (run #2) with a deprotonated E311 (TM7 Glu of PepT_Sh_) to stabilize a salt bridge between E311 and R44 (TM1 Arg of PepT_Sh_) and, as predicted, the salt bridge remained stable in the simulations and the transporter populated the IF state (**Fig. 5A**, blue histogram; **Fig. S5**, left panel). Notably, the system also samples an inward-facing occluded (IF-OC) state featuring a partially closed intracellular gate in our 3.5-*μ*s simulation (**Fig. 5BCD**), and we discovered that this state is metastable since the transporter returns to the IF state after ∼ 400 ns in the IF-OC state (**Fig. 5C**). The biological relevance of the IF-OC state will be discussed below.

**FIGURE 5.**
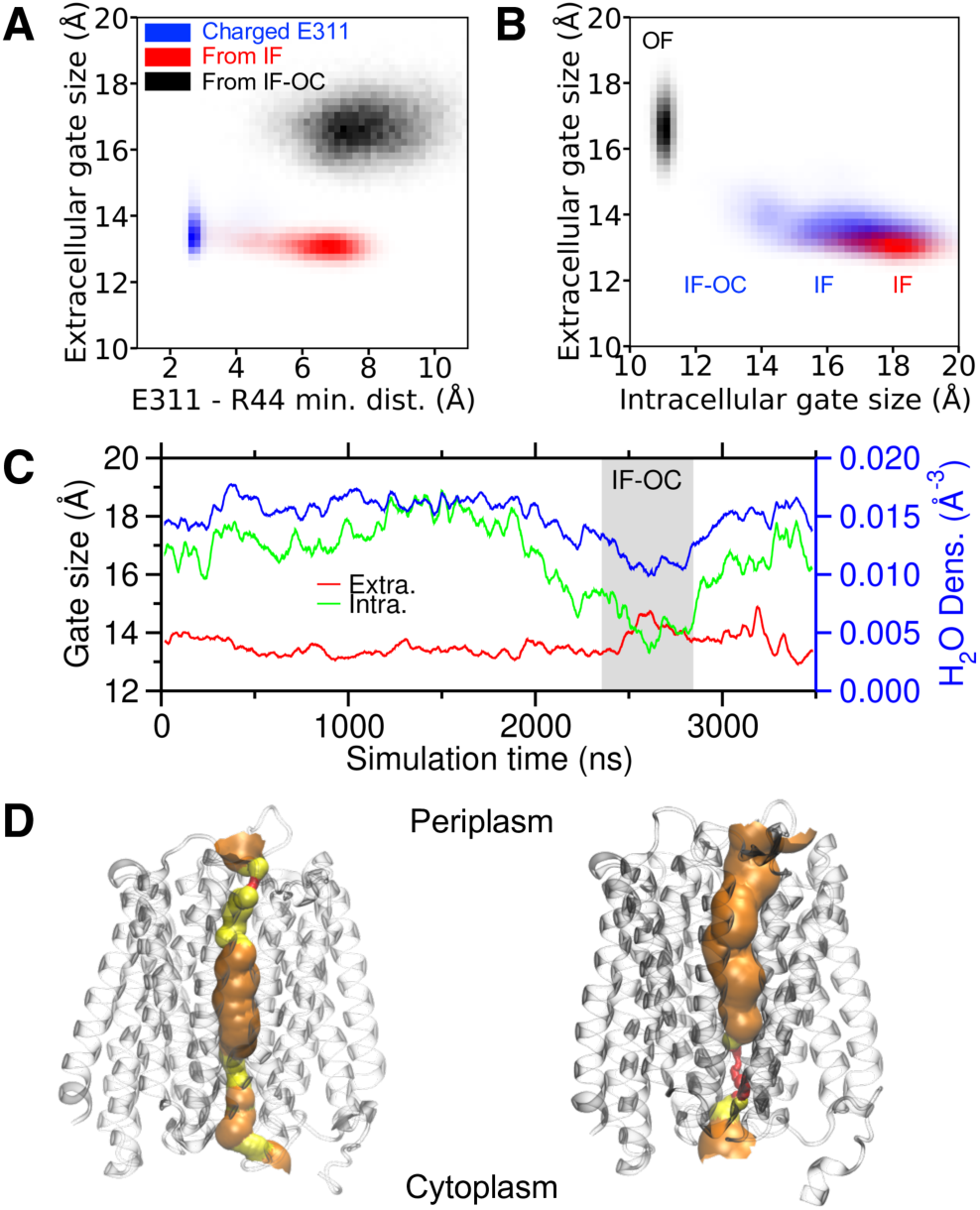
Proton-induced conformational change. **(A)** Two-dimensional histogram of the minimum distance between E311 and R44 heavy atoms, and the extracellular gate size of simulations E33^H^/E311^−^/E418^−^ (run #2-1, blue), E33^H^/E311^H^/E418^−^ initiated from an inward-facing occluded conformation (run #4-1, black), and E33^H^/E311^−^/E418^−^ initiated from an inward-facing conformation (run #5-1, red). **(B)** Two-dimensional histogram of the intracellular and extracellular gate sizes. **(C)** The gate sizes and the water density around the intracellular gate in the E33^H^/E311^−^/E418^−^ simulation (run #2-1). The region corresponding to the inward-facing occluded state is highlighted by grey. A running average with a 20-ns window was performed on the time series. **(D)** The pore radius profile of the MD-sampled inward-facing occluded state. **(E)** The pore radius profile of the MD-sampled outward-facing state.

To model the PT from the periplasm to E311, we initiated E33^H^/E311^H^/E418^−^ simulations (run #5) from equilibrated IF structures sampled by the E33^H^/E311^−^/E418^H^ simulations (run #2) and, as a result, the E311–R44 salt bridge broke and the two residues separated. Interestingly, in our 1-μs simulations of three independent runs, the transporter remained in the IF state (**Fig. 5AB**, red histogram; **Fig. S4**, bottom panels; **Fig. S5**, right panel), which is indeed consistent with prior MD investigations. This contradicts the idea that the protonation of the TM7 Glu opens the extracellular gate, since we found only a weak correlation between the TM7 Glu–TM1 Arg distance and the extracellular gate size, shown by a wide range of Glu-Arg distances correlating to a narrow gate size distribution (**Fig. 5A**, red histogram).

However, in E33^H^/E311^H^/E418^−^ simulations (run #4) initiated from the IF-OC state sampled by the E33^H^/E311^−^/E418^−^ simulations (run #2), the transporter undergoes a fast conformational change into the OF state (∼100 ns) following the acceptance of a proton by E311, as seen in two separate runs (#4-1 and #4-2; **Fig. S4**, middle panels; **Fig. S5**, middle panel). It is also interesting to note that in the third separate run the IF-OC state immediately switched back to the IF state (run #4-3; **Fig. S4**, middle panels), indicating some reversibility in the conformational transitions of this transporter, in the sense that IF-OC to IF transition is rare but possible even with a protonated E311. The slow shutting of the intracellular gate is thought to be the cause for the restriction of a straight transition from IF to OF state. Notably, there is a clear correlation between the intracellular gate size and the water density between helix tips of TM4, 5, 10, and 11 that form the intracellular gate (**Fig. 5C**, blue line), indicating that fluctuations in hydration may be important for the intracellular gate closure. The collective motion of water molecules can represent a slow degree of freedom, and an enhanced conformational sampling that omits it may suffer from strong hysteresis and produce inaccurate free energetics (see **Fig. S6** as an example of slow convergence of conformational free energies). Interestingly, when starting from the equilibrated OF structures, deprotonation of E311 does not result in the closing of the extracellular gate in our three independent E33^H^/E311^−^/E418^−^ simulations (run #6; **Fig. S7**), implying that the closing of the gate is as slow as the intracellular gate, which may be the rate-limiting step of the entire functional cycle. This behavior is in contrast with a kinetic model (71,72) in which multiple stochastic paths are possible; in this case the closing of the gate may provide a kinetic funnel bottleneck through which all paths are focused.

The equilibrated OF conformation was superimposed with a recently resolved OF structure of rat PepT2 (PDB: 7NQK) (73) to determine if the structure was a simulation artifact. The two structures were aligned using a maximum likelihood approach (74,75) and they overlap extremely well (*C*^*α*^ maximum likelihood RMSD = 0.95 Å), as illustrated in **Fig. 6**, and a deeper inspection of the critical residues TM 7 and 10 glutamates/aspartates and TM1 arginine/lysine, as well as the well-conserved ExxER/K motif, reveals a significant overlap between those two structures. These observations support that the OF state sampled from our MD simulations are a physical state of the transporter. As derived from the findings presented above, we directly observe that the titration of TM7 E311 initiates the opening of the extracellular gate via the breaking of the E311–R44 salt bridge, and that the underlying mechanism is more complex than previously anticipated. Although a protonated E311 loses the salt bridge with R44, potentially allowing the extracellular gate structural flexibility, this is insufficient for the transporter to directly switch from an IF to an OF state. The rate-limiting step for the transition is to find an IF-OC state and potentially the coupled hydration fluctuation that may take microseconds or more. Once the IF-OC state has been reached, the transition to the OF state occurs quickly, as observed in runs #4-1 and #4-2, in roughly 100 nanoseconds.

**FIGURE 6.**
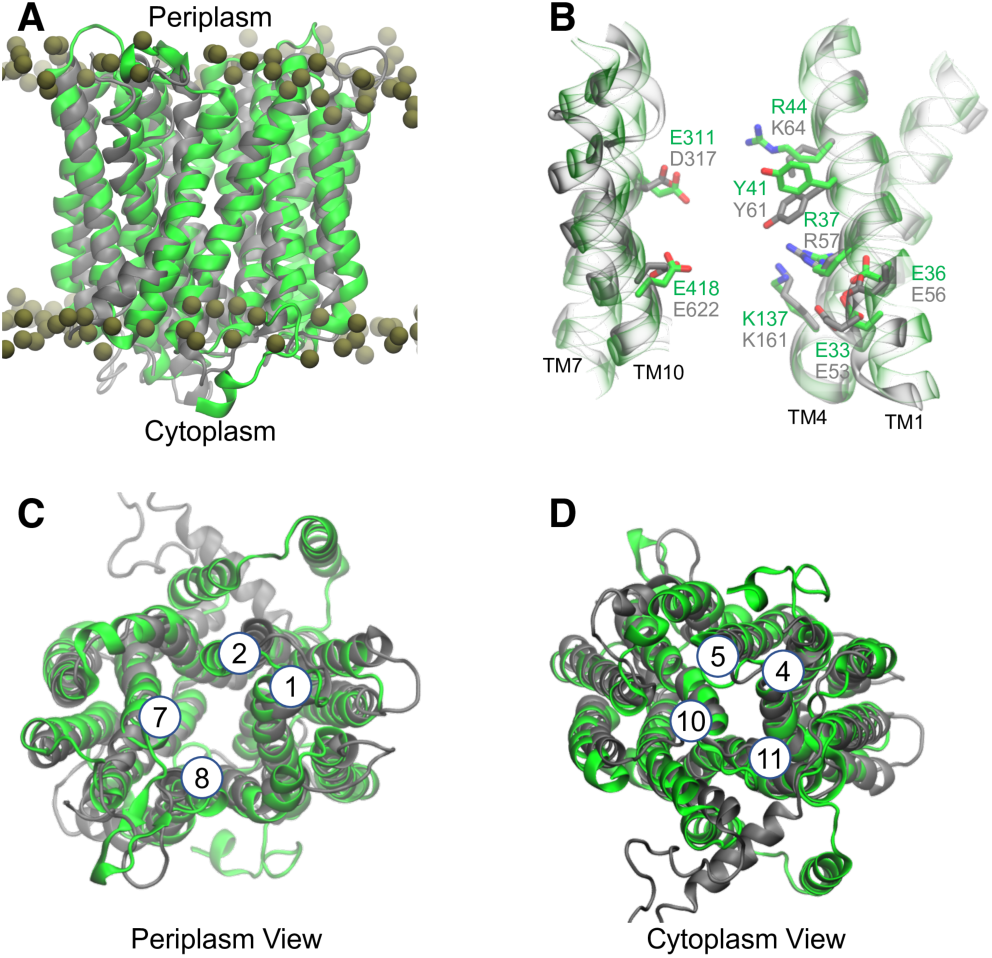
Structural comparison between PepT_Sh_ and PepT2. **(A)** Superposition of the MD-sampled outward-facing PepT_Sh_ (green) with a cryo-EM outward-facing PepT2 (grey; PDB: 7NQK). The lipid phosphorus atoms are represented by dark yellow spheres. Comparison of **(B)** key residue, **(C)** extracellular gate and **(D)** intracellular gate between PepT_Sh_ and PepT2. The helices that form the gates are labeled in **(C)** and **(D)**.

### Facile Proton Transfer between TM7 and TM10 via water

Given that TM7 E311 requires losing a proton to seal the extracellular gate while TM10 E418 requires a proton to deliver the ligand into the cytosol, it is a plausible assumption that the proton needs to be transported from E311 to E418. According to the results we showed previously, the proton can drive the conformational change and ligand transport, and now we want to explore how the conformation and ligand, in turn, effect this PT. Toward that goal, we implemented the MS-RMD methodology to simulate explicit proton transport for an all-atom reactive potential energy surface (proton transporting via the Grotthuss shuttle mechanism) in both the IF and OF states, as well as the apo and holo forms, to calculate the free energetics of this PT step with these coupled motions. Interestingly, as seen in **Fig. 7A**, the two potentials of mean force (PMF; free energy profiles) of PT in apo OF and apo IF differ only slightly. The difference in the free energy well corresponding to a protonated E311 reveals that the proton on E311 in the IF state is about 1 kcal/mol less stable than that in OF, which is a result of a closer positively charged R44 in the IF state.

**FIGURE 7.**
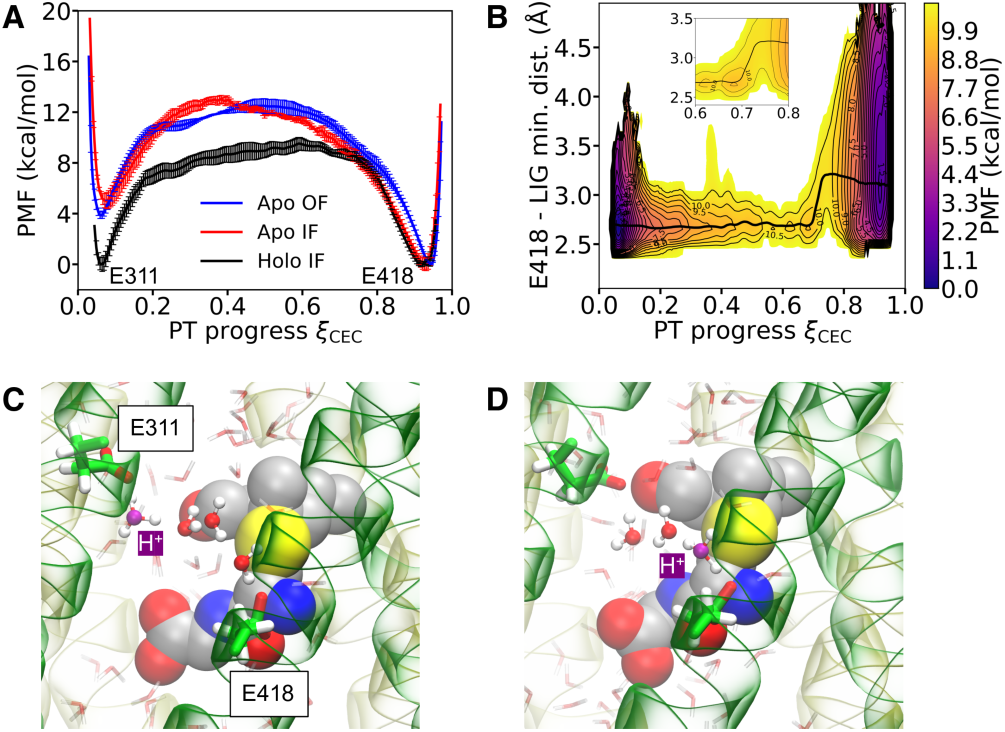
Characterization of proton transport between TM7 and TM10. **(A)** Potential of mean force (free energy profile) for proton transport between E311 and E418. **(B)** Two-dimensional potential of mean force of proton transport between E311 and E418 with the minimum distance between E418 carboxyl and ligand N-terminal nitrogen atoms. The inset is a zoom-in showing the strongly coupled region between proton and ligand. **(C) & (D)** Molecular figures showing Grotthuss proton shuttling mechanism when the ligand is present. The most probable hydronium oxygen is highlighted in purple. The ligand is shown in the van-der-Waals representation. The N-terminal bundle of the protein is shown in transparent yellow and the C-terminal bundle is shown in transparent green.

Another notable variation in PT behavior is the location of the free energy barrier maximum, which is closer to E311 in apo IF than in apo OF. In the IF state, the closure of the extracellular gate restricts hydration around E311, resulting in less hydration (solvation) for the excess proton (H^+^) in that region and as a result, an earlier barrier is seen in the PMF. In spite of these differences, both PMFs reveal that the proton is more thermodynamically stable when bonded to E418 than E311, showing an E311→ E418 proton movement, and this preference will be even more significant when an inward proton gradient across the membrane is taken into account.

In addition to the thermodynamic preference, the calculated rate constants (**Table 1**) confirm that the PT is facile in both apo conformational states. When comparing to the turnover rate of the protein *in vitro* ∼ 2 seconds (19), this PT will not be rate-limiting in the transporter functional cycle.

**Table 1.** Calculated proton transport rates between TM7 E311 and TM10 E418.

|  |  |  |  |
| --- | --- | --- | --- |
| System | Apo IF | Apo OF | Holo IF |
| $k_{E311 \rightarrow E418} (\mu s^{-1})$ | $19.2 \pm 0.8$ | $1.6 \pm 0.4$ | $0.54 \pm 0.26$ |
| $k_{E418 \rightarrow E311} (\mu s^{-1})$ | $1.5 \pm 1.4 \times 10^{-2}$ | $3.8 \pm 1.6 \times 10^{-3}$ | $0.41 \pm 0.39$ |

As mentioned in the preceding section, the TM10 E418 is correlated to ligand transport, and its protonation results in cytosolic ligand release. Here, we again used the MS-RMD method in conjunction with enhanced sampling to investigate how the proton could be transferred from E311 to E418 in the presence of the ligand. The presence of ligand does not significantly alter the overall shape of the PT free energy, as seen in **Fig. 7A**, but it does affect the relative stability between protonated E311 and E418. Because of the energy gained through forming a salt bridge between deprotonated E418 and the ligand N-terminus, the excess proton is now equally stable on both glutamates, and the PT direction will be fully determined by the direction of the proton gradient. Interestingly, the bulky sidechain does not fully remove water between the glutamates, allowing an excess proton to travel through connected water wires via Grotthuss shuttling (76,77) (**Fig. 7CD**). As a result, the PT rate is slower than in the apo case (**Table 1**), but it is still a feasible PT and would not constitute the rate-limiting step of the entire cycle. This facile PT mechanism entails a “decoupling” between proton and ligand transport in the sense that the ligand is not needed to participate and has only a modest impact on proton movement, thus adding to our understanding of why the transporter may use the proton gradient to transport a wide range of ligands. The joint free energy profile of PT progress with the E418–ligand distance (**Fig. 7B**), on the other hand, depicts the coupling between proton and ligand. When a proton is bound to E418, the PMF shows more flexibility in the direction of E418–ligand distance, consistent with the observations in classical MD that protonation of E418 eventually triggers ligand release. At equilibrium, the system motion tends to follow the minimum free energy path (MFEP), and the ramped slope of the MFEP, especially in the range of 0.6 < *ξ*_CEC_ < 0.8 (**Fig. 7B**, inset), reveals that the motion of the proton can drive the motion of the ligand and that the ligand can also be the driving force of the proton.

## DISCUSSION

POT family proteins represent secondary active transporters that make use of the proton electrochemical gradient to transport various peptides and their analogs into cells. Unraveling the mechanism of how transport is coupled to the proton gradient is important for understanding and improving drug transport, pharmacokinetics, and oral bioavailability via this SLC system. However, it remains challenging to experimentally track the molecular motions in real-time at an atomistic resolution, thus limiting a direct experimental examination of the detailed transport process and discovery of the important interactions. MD simulations, on the other hand, are a valuable tool for studying complicated molecular systems and processes to enable direct insight into atomic motions as well as the calculations of associated free energetics and kinetics.

In this study and building upon our prior structural and biochemical characterizations, we used comprehensive classical and reactive MD along with enhanced free energy sampling to elucidate the proton coupling mechanism of PepT_Sh_, a member of the POT family. As summarized in **Fig. 8**, based on our results we suggest a schematic functional cycle where the proton flow mediated by the transporter drives the conformational switching and ligand movement via altering the charge states of the TM7 and TM10 glutamates. However, we should emphasize that proteins are stochastic molecular machines and the hopping between the microstates shown in **Fig. 8** must be considered to be probabilistic. Moreover, there may exist multiple possible transition pathways connecting the states as hypothesized for other transporters, such as ClC-ec1 (71,72) and PiPT (78). As such, it should also be the case in PepT_Sh_, and for example, in one pathway (**Fig. 8**, B→F→E→J), that one single proton reaches TM10 E418 after ligand binding and activates its release. Alternatively, the proton may possibly reach E418 before the ligand binds, and the ligand tail inserts into the TM7-TM10 pocket, resulting in the state captured in the crystal structure (PDB: 6EXS) (**Fig. 8**, B→C→G). The proton then dissociates from E418 and enters the cytosol (**Fig. 8**, G→L), forming an E418–ligand salt bridge (**Fig. 8**, L→K), which is followed by another proton binding to E418 and releasing the ligand (**Fig. 8**, K→F→E→J). We note that in the case of ClC-ec1, the rate-limiting step is not completely restricted to a specific elementary process but is more diverse among multiple pathways, while here in this peptide transporter it may be always the slow global conformational change that dominates over all the various pathway rates. In **Fig. 8**, we mainly outlined the primary functioning pathways that best represent the data at hand, but a more complete quantitative description of the whole functional cycle may be required to fully understand the coupling between protein, ligand, and proton motions, as well as their pH dependence and stoichiometry. To achieve that, even more extensive and converged conformational enhanced sampling of the protein is necessary to accurately estimate the transition energy and rates.

**FIGURE 8.**
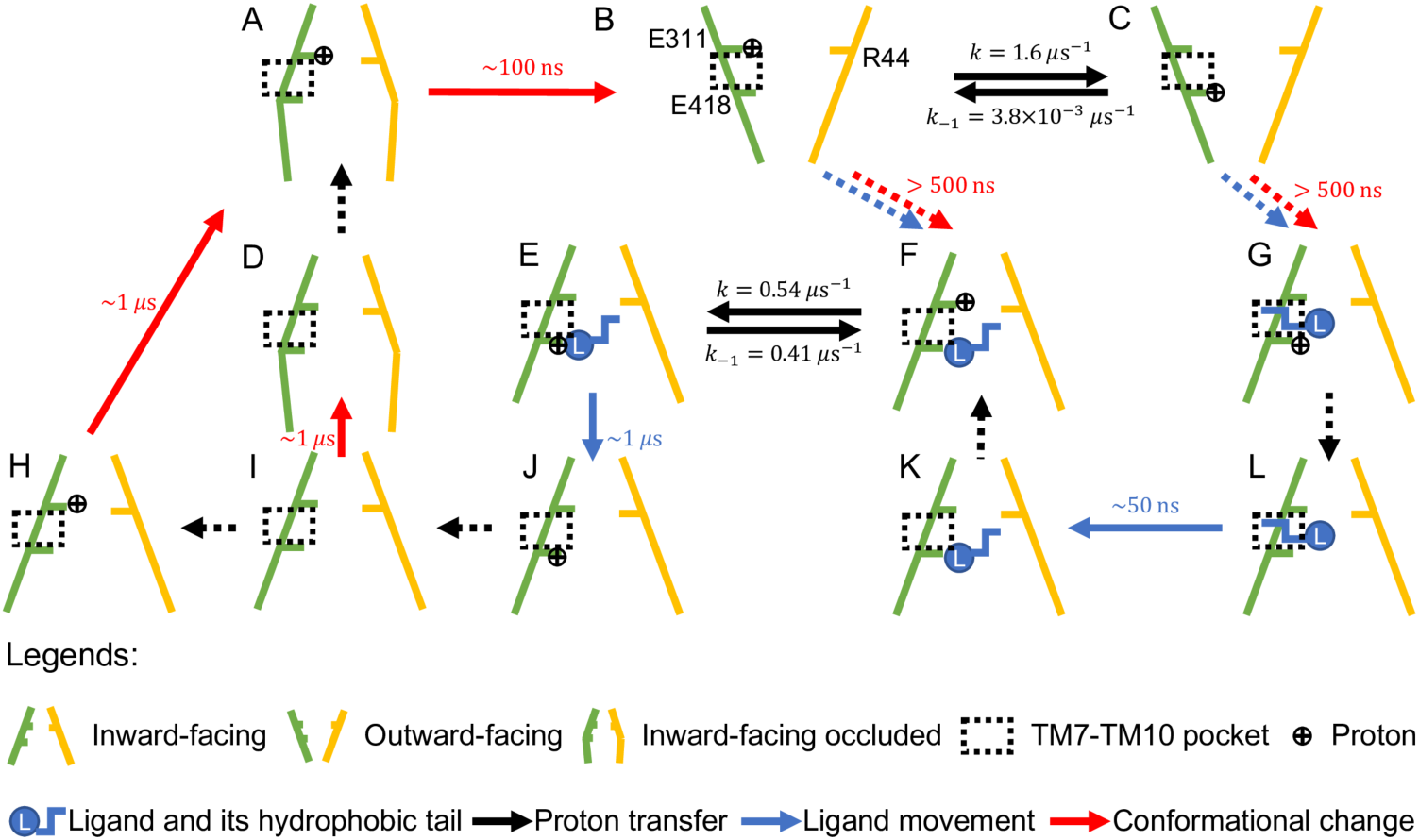
Schematic diagram of the transporter functional conformational and kinetic cycle. The dashed arrows represent the transitions whose reaction rates are not known yet from the performed simulations. The N-terminal bundle is represented by yellow sticks and the C-terminal bundle is colored in green. Note the PT rates were computed without a pH gradient.

Because hydration may also play a critical and entangled role in conformational changes, a reaction coordinate describing it should be properly defined and employed in the free energy sampling. However, the inclusion of additional degrees of freedom in such enhanced sampling may result in an excessive computing overhead when dealing with a high-dimensional CV space. In this scenario, machine learning and statistical approaches (79-82) could effectively reduce the sampling dimensionality and intelligently select the most significant phase space region upon which to focus.

We note that our discussions up to now provides only a limited understanding of the critical importance of the ExxER/K motif. In comparison with only IF and IF-OC states observed in the protonated E33 simulations (**Fig. 5C**), our E33^−^/E311^−^/E418^−^ simulations (run #7) show that the intracellular gate becomes more flexible with a deprotonated E33, as the IF-OC (run #7-1; **Fig. S8**, top panel), IF (run #7-2; **Fig. S8**, middle panel), and OC (run #7-3; **Fig. S8**, bottom panel) states were sampled, but a more rigorous conformational free energy landscape is needed to draw a firm conclusion. In the current simulations, the charged E33 serves as an attraction to stabilize the unusual interaction between R37 and K137 observed in PepT_Sh_ as well as in the cryo-EM rat PepT2 structure (73). We further note that the current force field-based description may be inaccurate for a possible proton sharing between R37 and K137, but a quantum mechanical treatment of that motif would be too computationally expensive to achieve adequate conformational sampling. In future work, it may be possible to train an effective potential from highly accurate electronic structure calculations that corrects the conventional force fields for better modeling this novel Arg-Lys interaction, in light of the recent developments deep learning potentials using *ab initio* calculations (83,84).

## SUPPORTING MATERIAL

Supporting material can be found online at https://doi.org/10.1016/j.bpj.xxxx.xx.xxx.

## AUTHOR CONTRIBUTIONS

G.A.V. and S.N. conceptualized and supervised the project. C.L. and Z.Y. performed and analyzed all MD simulations. C.L., Z.Y., S.N., and G.A.V. wrote the manuscript.

## ACKNOWLEDGMENTS

This research was supported by the National Institute of General Medical Sciences (NIGMS) of the U.S. National Institutes of Health (NIH) through grant R01 GM053148. Computational resources were provided by the Research Computing Center (RCC) at the University of Chicago.

## REFERENCES

1. Daniel, H., and G. Kottra. 2004. The proton oligopeptide cotransporter family SLC15 in physiology and pharmacology. Pflügers Arch. 447:610–618.

2. Smith, D. E., B. Clémençon, and M. A. Hediger. 2013. Proton-coupled oligopeptide transporter family SLC15: Physiological, pharmacological and pathological implications. Mol. Asp. Med. 34:323–336.

3. Rubio-Aliaga, I., and H. Daniel. 2002. Mammalian peptide transporters as targets for drug delivery. Trends Pharmacol. Sci. 23:434–440.

4. International Transporter Consortium. 2010. Membrane transporters in drug development. Nat. Rev. Drug Discov. 9:215–236.

5. Lin, L., S. W. Yee, R. B. Kim, and K. M. Giacomini. 2015. SLC transporters as therapeutic targets: emerging opportunities. Nat. Rev. Drug Discov. 14:543–560.

6. Solcan, N., J. Kwok, P. W. Fowler, A. D. Cameron, D. Drew, S. Iwata, and S. Newstead. 2012. Alternating access mechanism in the POT family of oligopeptide transporters. EMBO J. 31:3411–3421.

7. Newstead, S., D. Drew, A. D. Cameron, V. L. G. Postis, X. B. Xia, P. W. Fowler, J. C. Ingram, E. P. Carpenter, M. S. P. Sansom, M. J. McPherson, S. A. Baldwin, and S. Iwata. 2011. Crystal structure of a prokaryotic homologue of the mammalian oligopeptide–proton symporters, PepT1 and PepT2. EMBO J. 30:417–426.

8. Quistgaard, E. M., C. Löw, F. Guettou, and P. Nordlund. 2016. Understanding transport by the major facilitator superfamily (MFS): structures pave the way. Nat. Rev. Mol. Cell Biol. 17:123–132.

9. Drew, D., and O. Boudker. 2016. Shared Molecular Mechanisms of Membrane Transporters. Annu. Rev. Biochem. 85:543–572.

10. Newstead, S. 2017. Recent advances in understanding proton coupled peptide transport via the POT family. Curr. Opin. Struct. Biol. 45:17–24.

11. Parker, J. L., C. Li, A. Brinth, Z. Wang, L. Vogeley, N. Solcan, G. Ledderboge-Vucinic, J. M. J. Swanson, M. Caffrey, G. A. Voth, and S. Newstead. 2017. Proton movement and coupling in the POT family of peptide transporters. Proc. Natl. Acad. Sci. U.S.A. 114:13182–13187.

12. Huang, Y. F., M. J. Lemieux, J. M. Song, M. Auer, and D. N. Wang. 2003. Structure and Mechanism of the Glycerol-3-Phosphate Transporter from Escherichia coli. Science 301:616–620.

13. Newstead, S. 2015. Molecular insights into proton coupled peptide transport in the PTR family of oligopeptide transporters. Biochim. Biophys. Acta 1850:488–499.

14. Doki, S., H. E. Kato, N. Solcan, M. Iwaki, M. Koyama, M. Hattori, N. Iwase, T. Tsukazaki, Y. Sugita, H. Kandori, S. Newstead, R. Ishitani, and O. Nureki. 2013. Structural basis for dynamic mechanism of proton-coupled symport by the peptide transporter POT. Proc. Natl. Acad. Sci. U.S.A. 110:11343–11348.

15. Immadisetty, K., J. Hettige, and M. Moradi. 2017. What Can and Cannot Be Learned from Molecular Dynamics Simulations of Bacterial Proton-Coupled Oligopeptide Transporter GkPOT? J. Phys. Chem. B 121:3644–3656.

16. Selvam, B., S. Mittal, and D. Shukla. 2018. Free Energy Landscape of the Complete Transport Cycle in a Key Bacterial Transporter. ACS Cent. Sci. 4:1146–1154.

17. Batista, M. R. B., A. Watts, and A. J. Costa-Filho. 2019. Exploring Conformational Transitions and Free-Energy Profiles of Proton-Coupled Oligopeptide Transporters. J. Chem. Theory Comput. 15:6433–6443.

18. Aduri, N. G., M. Montefiori, R. Khalil, M. Gajhede, F. S. Jørgensen, and O. Mirza. 2019. Molecular Dynamics Simulations Reveal the Proton:Peptide Coupling Mechanism in the Bacterial Proton-Coupled Oligopeptide Transporter YbgH. ACS Omega 4:2040–2046.

19. Minhas, G. S., D. Bawdon, R. Herman, M. Rudden, A. P. Stone, A. G. James, G. H. Thomas, and S. Newstead. 2018. Structural basis of malodour precursor transport in the human axilla. eLife 7:e34995.

20. Collins, K. J. 1989. Sweat Glands: Eccrine and Apocrine. In Pharmacology of the Skin I, 1st ed.; M. W. Greaves and S. Shuster, Eds. Springer-Verlag, Berlin, Heidelberg, Germany; pp. 193–212.

21. Minhas, G. S., and S. Newstead. 2019. Structural basis for prodrug recognition by the SLC15 family of proton-coupled peptide transporters. Proc. Natl. Acad. Sci. U.S.A. 116:804–809.

22. Minhas, G. S., and S. Newstead. 2020. Recent advances in understanding prodrug transport through the SLC15 family of proton-coupled transporters. Biochem. Soc. Trans. 48:337–346.

23. Wu, Y., H. Chen, F. Wang, F. Paesani, and G. A. Voth. 2008. An Improved Multistate Empirical Valence Bond Model for Aqueous Proton Solvation and Transport. J. Phys. Chem. B 112:467–482.

24. Lee, S., R. Liang, G. A. Voth, and J. M. J. Swanson. 2016. Computationally Efficient Multiscale Reactive Molecular Dynamics to Describe Amino Acid Deprotonation in Proteins. J. Chem. Theory Comput. 12:879– 891.

25. Jo, S., T. Kim, and W. Im. 2007. Automated Builder and Database of Protein/Membrane Complexes for Molecular Dynamics Simulations. PLoS ONE 2:e880.

26. Jo, S., T. Kim, V. G. Iyer, and W. Im. 2008. CHARNIM-GUI: A Web-Based Grraphical User Interface for CHARMM. J. Comput. Chem. 29:1859–1865.

27. Durell, S. R., B. R. Brooks, and A. Ben-Naim. 1994. Solvent-Induced Forces between Two Hydrophilic Groups. J. Phys. Chem. 98:2198–2202.

28. MacKerell Jr., A. D., D. Bashford, M. Bellott, R. L. Dunbrack Jr., J. D. Evanseck, M. J. Field, S. Fischer, J. Gao, H. Guo, S. Ha, D. Joseph-McCarthy, L. Kuchnir, K. Kuczera, F. T. Lau, C. Mattos, S. Michnick, T. Ngo, D. T. Nguyen, B. Prodhom, W. E. Reiher, B. Roux, M. Schlenkrich, J. C. Smith, R. Stote, J. Straub, M. Watanabe, J. Wiórkiewicz-Kuczera, D. Yin, and M. Karplus. 1998. All-Atom Empirical Potential for Molecular Modeling and Dynamics Studies of Proteins. J. Phys. Chem. B 102:3586–3616.

29. MacKerell Jr., A. D., M. Feig, and C. L. Brooks III. 2004. Improved Treatment of the Protein Backbone in Empirical Force Fields. J. Am. Chem. Soc. 126:698–699.

30. Mackerell Jr., A. D., M. Feig, and C. L. Brooks III. 2004. Extending the Treatment of Backbone Energetics in Protein Force Fields: Limitations of Gas-Phase Quantum Mechanics in Reproducing Protein Conformational Distributions in Molecular Dynamics Simulations. J. Comput. Chem. 25:1400–1415.

31. Klauda, J. B., R. M. Venable, J. A. Freites, J. W. O’Connor, D. J. Tobias, C. Mondragon-Ramirez, I. V. Vorobyov, A. D. MacKerell Jr., and R. W. Pastor. 2010. Update of the CHARMM All-Atom Additive Force Field for Lipids: Validation on Six Lipid Types. J. Phys. Chem. B 114:7830–7843.

32. Vanommeslaeghe, K., E. Hatcher, C. Acharya, S. Kundu, S. Zhong, J. Shim, E. Darian, O. Guvench, P. Lopes Vorobyov, and A. D. Mackerell Jr. 2010. CHARMM general force field: A force field for drug-like molecules compatible with the CHARMM all-atom additive biological force fields. J. Comput. Chem. 31:671–690.

33. Darden, T., D. York, and L. Pedersen. 1993. Particle mesh Ewald: An N•log(N) method for Ewald sums in large systems. J. Chem. Phys. 98:10089–10092.

34. Essmann, U., L. Perera, M. L. Berkowitz, T. Darden, H. Lee, and L. G. Pedersen. 1995. A smooth particle mesh Ewald method. J. Chem. Phys. 103:8577–8593.

35. Wu, E. L., X. Cheng, S. Jo, H. Rui, K. C. Song, E. M. Dávila-Contreras, Y. Qi, J. Lee, V. Monje-Galvan, R. M. Venable, J. B. Klauda, and W. Im. 2014. CHARMM-GUI Membrane Builder Toward Realistic Biological Membrane Simulations. J. Comput. Chem. 35:1997–2004.

36. Hess, B., H. Bekker, H. J. C. Berendsen, and J. G. E. M. Fraaije. 1997. LINCS: A Linear Constraint Solver for Molecular Simulations. J. Comput. Chem. 18:1463–1472.

37. Nosé, S. 1984. A molecular dynamics method for simulations in the canonical ensemble. Mol. Phys. 52:255–268.

38. Hoover, W. G. 1985. Canonical dynamics: Equilibrium phase-space distributions. Phys. Rev. A 31:1695–1697.

39. Parrinello, M., and A. Rahman. 1981. Polymorphic transitions in single crystals: A new molecular dynamics method. J. Appl. Phys. 52:7182–7190.

40. Abraham, M. J., T. Murtola, R. Schulz, S. Páll, J. C. Smith, B. Hess, and E. Lindahl. 2015. GROMACS: High performance molecular simulations through multi-level parallelism from laptops to supercomputers. SoftwareX 1-2:19–25.

41. Li, C., and G. A. Voth. 2021. Accurate and Transferable Reactive Molecular Dynamics Models from Constrained Density Functional Theory. J. Phys. Chem. B 125:10471–10480.

42. Hockney, R. W., and J. W. Eastwood. 1981. Particle-Particle–Particle-Mesh (P3M) Algorithms. In Computer Simulation Using Particles, 1st ed.; R. W. Hockney and J. W. Eastwood, Eds. McGraw-Hill International Book Co., New York, NY, USA; pp. 267–304.

43. Plimpton, S. 1995. Fast Parallel Algorithms for Short-Range Molecular Dynamics. J. Comput. Phys. 117:1– 19.

44. Yamashita, T., Y. X. Peng, C. Knight, and G. A. Voth. 2012. Computationally Efficient Multiconfigurational Reactive Molecular Dynamics. J. Chem. Theory Comput. 8:4863–4875.

45. Barducci, A., G. Bussi, and M. Parrinello. 2008. Well-Tempered Metadynamics: A Smoothly Converging and Tunable Free-Energy Method. Phys. Rev. Lett. 100:020603.

46. Cuma, M., U. W. Schmitt, and G. A. Voth. 2001. A multi-state empirical valence bond model for weak acid dissociation in aqueous solution. J. Phys. Chem. A 105:2814–2823.

47. Torrie, G. M., and J. P. Valleau. 1977. Non-Physical Sampling Distributions in Monte-Carlo Free-Energy Estimation - Umbrella Sampling. J. Comput. Phys. 23:187–199.

48. Tribello, G. A., M. Bonomi, D. Branduardi, C. Camilloni, and G. Bussi. 2014. PLUMED 2: New feathers for an old bird. Comput. Phys. Commun. 185:604–613.

49. Rosta, E., and G. Hummer. 2015. Free Energies from Dynamic Weighted Histogram Analysis Using Unbiased Markov State Model. J. Chem. Theory Comput. 11:276–285.

50. Sicard, F., V. Koskin, A. Annibale, and E. Rosta. 2021. Position-Dependent Diffusion from Biased Simulations and Markov State Model Analysis. J. Chem. Theory Comput. 17:2022–2033.

51. Li, C., and G. A. Voth. 2021. A quantitative paradigm for water-assisted proton transport through proteins and other confined spaces. Proc. Natl. Acad. Sci. U.S.A. 118:e2113141118.

52. Szabo, A., K. Schulten, and Z. Schulten. 1980. First passage time approach to diffusion controlled reactions. J. Chem. Phys. 72:4350–4357.

53. Chen, W., Y. Huang, and J. Shen. 2016. Conformational Activation of a Transmembrane Proton Channel from Constant pH Molecular Dynamics. J. Phys. Chem. Lett. 7:3961–3966.

54. Wallace, J. A., and J. K. Shen. 2011. Continuous Constant pH Molecular Dynamics in Explicit Solvent with pH-Based Replica Exchange. J. Chem. Theory Comput. 7:2617–2629.

55. Brooks, B. R., C. L. Brooks III, A. D. Mackerell, L. Nilsson, R. J. Petrella, B. Roux, Y. Won, G. Archontis, C. Bartels, S. Boresch, A. Caflisch, L. Caves, Q. Cui, A. R. Dinner, M. Feig, S. Fischer, J. Gao, M. Hodoscek, W. Im, K. Kuczera, T. Lazaridis, J. Ma, V. Ovchinnikov, E. Paci, R. W. Pastor, C. B. Post, J. Z. Pu, M. Schaefer, B. Tidor, R. M. Venable, H. L. Woodcock, X. Wu, W. Yang, D. M. York, and M. Karplus. 2009. CHARMM: The Biomolecular Simulation Program. J. Comput. Chem. 30:1545–1614.

56. Lee, M. S., F. R. Salsbury Jr., and C. L. Brooks III. 2004. Constant-pH Molecular Dynamics Using Continuous Titration Coordinates. Proteins 56:738–752.

57. Khandogin, J., and C. L. Brooks III. 2005. Constant pH Molecular Dynamics with Proton Tautomerism. Biophys. J. 89:141–157.

58. Steinbach, P. J., and B. R. Brooks. 1994. New Spherical-Cutoff Methods for Long-Range Forces in Macromolecular Simulation. J. Comput. Chem. 15:667–683.

59. Ryckaert, J.-P., G. Ciccotti, and H. J. C. Berendsen. 1977. Numerical Integration of the Cartesian Equations of Motion of a System with Constraints: Molecular Dynamics of n-Alkanes. J. Comput. Phys. 23:327–341.

60. Feller, S. E., Y. Zhang, R. W. Pastor, and B. R. Brooks. 1995. Constant pressure molecular dynamics simulation: The Langevin piston method. J. Chem. Phys. 103:4613–4621.

61. Loncharich, R. J., B. R. Brooks, and R. W. Pastor. 1992. Langevin Dynamics of Peptides: The Frictional Dependence of Isomerization Rates of N-Acetylalanyl-N’-Methylamide. Biopolymers 32:523–535.

62. Hockney, R. W. 1970. The Potential Calculation and Some Applications. Methods Comput. Phys. 9:135–211.

63. Im, W., M. S. Lee, and C. L. Brooks III. 2003. Generalized Born Model with a Simple Smoothing Function. J. Comput. Chem. 24:1691–1702.

64. Im, W., M. Feig, and C. L. Brooks III 2003. An Implicit Membrane Generalized Born Theory for the Study of Structure, Stability, and Interactions of Membrane Proteins. Biophys. J. 85:2900–2918.

65. Chen, J., W. Im, and C. L. Brooks III. 2006. Balancing Solvation and Intramolecular Interactions: Toward a Consistent Generalized Born Force Field. J. Am. Chem. Soc. 128:3728–3736.

66. Srinivasan, J., M. W. Trevathan, P. Beroza, and D. A. Case. 1999. Application of a pairwise generalized Born model to proteins and nucleic acids: inclusion of salt effects. Theor. Chem. Acc. 101:426–434.

67. Metropolis, N., and S. Ulam. 1949. The Monte Carlo Method. J. Am. Stat. Assoc. 44:335–341.

68. Hill, A. V. 1910. The possible effects of the aggregation of the molecules of haemoglobin on its dissociation curves. J. Physiol. 40:iv–vii.

69. Xu, L. Y., I. S. Haworth, A. A. Kulkarni, M. B. Bolger, and D. L. Davies. 2009. Mutagenesis and Cysteine Scanning of Transmembrane Domain 10 of the Human Dipeptide Transporter. Pharm. Res. 26:2358–2366.

70. Smart, O. S., J. G. Neduvelil, X. Wang, B. A. Wallace, and M. S. P. Sansom. 1996. HOLE: A program for the analysis of the pore dimensions of ion channel structural models. J. Mol. Graph. 14:354–360.

71. Mayes, H. B., S. Lee, A. D. White, G. A. Voth, and J. M. J. Swanson. 2018. Multiscale Kinetic Modeling Reveals an Ensemble of Cl–/H+ Exchange Pathways in ClC-ec1 Antiporter. J. Am. Chem. Soc. 140:1793–1804.

72. Yue, Z., A. Bernardi, C. Li, A. V. Mironenko, and J. M. J. Swanson. 2021. Toward a Multipathway Perspective: pH-Dependent Kinetic Selection of Competing Pathways and the Role of the Internal Glutamate in Cl–/H+ Antiporters. J. Phys. Chem. B 125:7975–7984.

73. Parker, J. L., J. C. Deme, Z. Y. Wu, G. Kuteyi, J. D. Huo, R. J. Owens, P. C. Biggin, S. M. Lea, and S. Newstead. 2021. Cryo-EM structure of PepT2 reveals structural basis for proton-coupled peptide and prodrug transport in mammals. Sci. Adv. 7:eabh3355.

74. Theobald, D. L., and D. S. Wuttke. 2008. Accurate Structural Correlations from Maximum Likelihood Superpositions. PLoS Comput. Biol. 4:e43.

75. Theobald, D. L., and P. A. Steindel. 2012. Optimal simultaneous superpositioning of multiple structures with missing data. Bioinformatics 28:1972–1979.

76. von Grotthuss, C. J. D. 1806. Sur la décomposition de l’eau et des corps qu’elle tient en dissolution à l’aide de l’électricité galvanique. Ann. Chim. 58:54–73.

77. Knight, C., and G. A. Voth. 2012. The Curious Case of the Hydrated Proton. Acc. Chem. Res. 45:101–109.

78. Liu, Y., C. Li, M. Gupta, N. Verma, A. K. Johri, R. M. Stroud, and G. A. Voth. 2021. Key computational findings reveal proton transfer as driving the functional cycle in the phosphate transporter PiPT. Proc. Natl. Acad. Sci. U.S.A. 118:e2101932118.

79. Sultan, M. M., and V. S. Pande. 2017. tICA-Metadynamics: Accelerating Metadynamics by Using Kinetically Selected Collective Variables. J. Chem. Theory Comput. 13:2440–2447.

80. McCarty, J., and M. Parrinello. 2017. A variational conformational dynamics approach to the selection of collective variables in metadynamics. J. Chem. Phys. 147:204109.

81. Mardt, A., L. Pasquali, H. Wu, and F. Noé. 2018. VAMPnets for deep learning of molecular kinetics. Nat. Commun. 9:5.

82. Zhang, L. F., H. Wang, and W. N. E. 2018. Reinforced dynamics for enhanced sampling in large atomic and molecular systems. J. Chem. Phys. 148:124113.

83. Zhang, L., J. Han, H. Wang, W. A. Saidi, R. Car, and E. Weinan. 2018. End-to-end Symmetry Preserving Interatomic Potential Energy Model for Finite and Extended Systems. In Proceedings of the 32nd International Conference on Neural Information Processing Systems, Montréal, Canada, 3–8 December, 2018; S. Bengio, H. Wallach, H. Larochelle, K. Grauman, N. Cesa-Bianchi, and R. Garnett, Eds. Curran Associates Inc., Red Hook, NY, USA; pp. 4441–4451.

84. Zhang, L. F., J. Q. Han, H. Wang, R. Car, and E. Weinan. 2018. Deep Potential Molecular Dynamics: A Scalable Model with the Accuracy of Quantum Mechanics. Phys. Rev. Lett. 120:143001.

