## Supplemental File for "Proton Coupling and the Multiscale Kinetic Mechanism of a Peptide Transporter"

### 1 Supplementary Figures and Tables

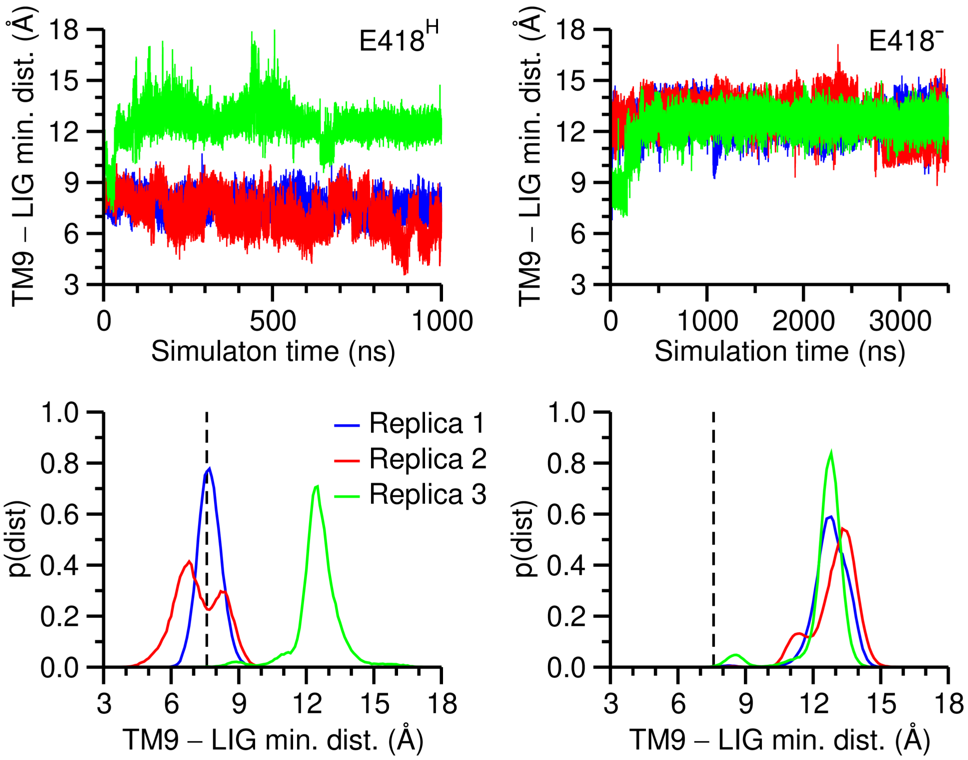


**Figure S1.** Time evolution (top panels) and distribution (bottom panels) of the minimum distance between the ligand and TM9 backbone atoms in E33$^{H}$/E311$^{-}$/E418$^{H}$ (run #1; left panels) and E33$^{H}$/E311$^{-}$/E418$^{-}$ (run #2; right panels) simulations. The value in the crystal structure was indicated by the dashed horizontal lin.

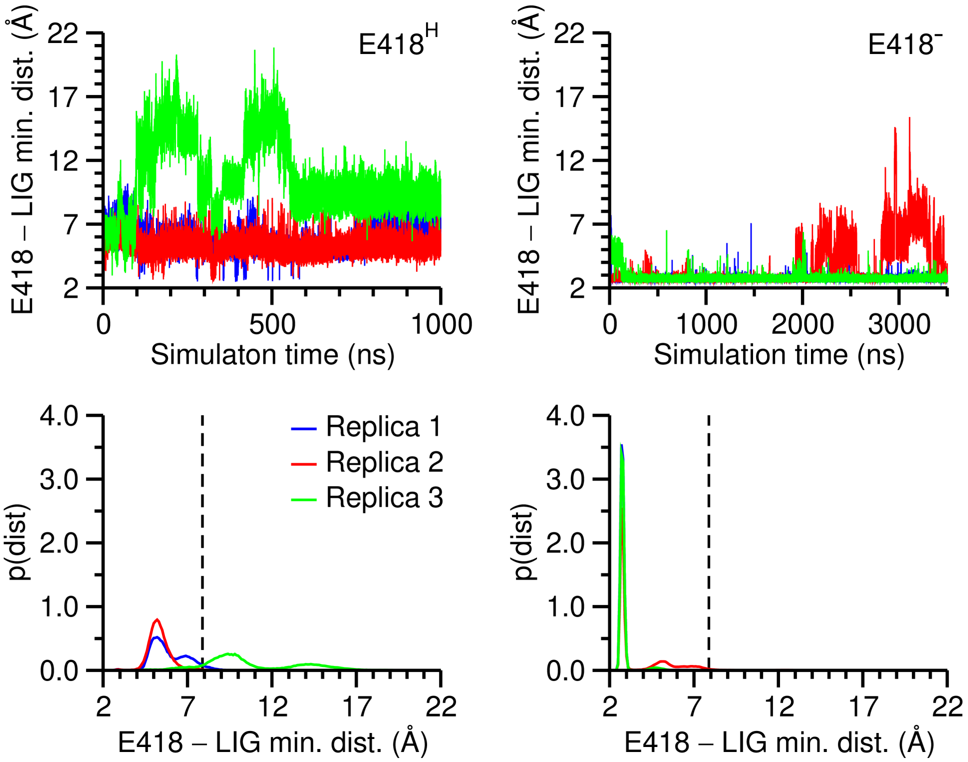


**Figure S2.** Time evolution (top panels) and distribution (bottom panels) of the minimum distance between the E418 carboxyl and the ligand N-terminus in $E33^{H}/E311^{-}/E418^{H}$ (run #1; left panels) and $E33^{H}/E311^{-}/E418^{-}$ (run #2; right panels) simulations. The value in the crystal structure was indicated by the dashed horizontal line.

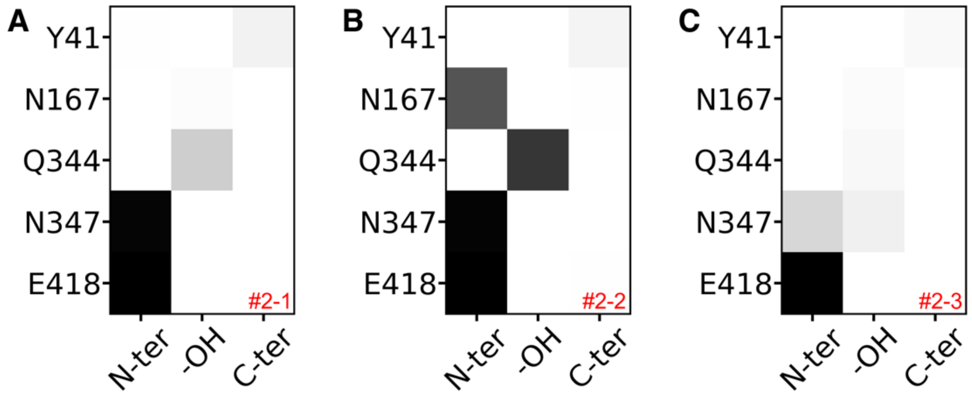


**Figure S3.** The contact map between ligand functional groups and binding-site residues in $E33^{H}/E311^{-}$ $/E418^{-}$ (run #2) simulations.

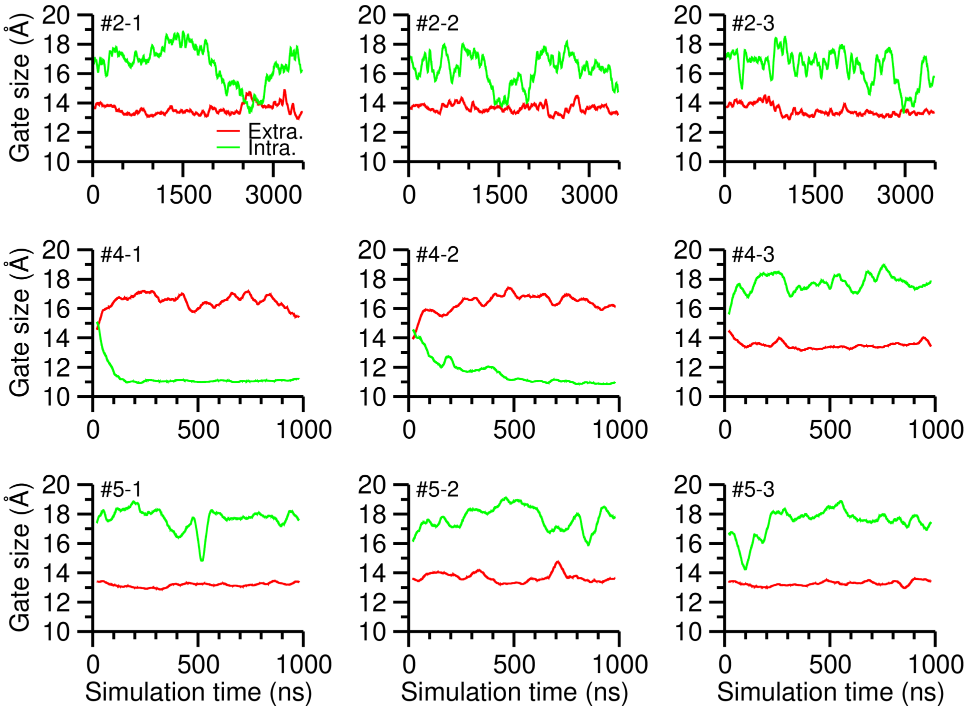


**Figure S4.** Time evolution of the extracellular (red lines) and intracellular (green lines) gate sizes in $E33^{H}/E311^{-}/E418^{-}$ (run #2), $E33^{H}/E311^{H}/E418^{-}$ (run #4), and $E33^{H}/E311^{H}/E418^{-}$ (run #5) simulations. Note runs #4 and #5 were initiated from the inward-facing occluded and inward-facing state, respectively. A running average with a 20-ns window was performed on the time series.

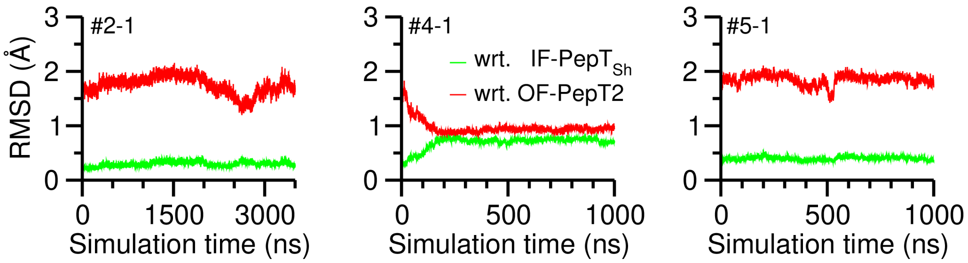


**Figure S5.** Time evolution of the maximum likelihood RMSD with respect to IF-$PepT_{Sh}$ (green lines; PDB: 6EXS) and OF-PepT2 (red lines; PDB: 7NQK) in simulations #2-1 ($E33^{H}/E311^{-}/E418^{-}$), #4-1 ($E33^{H}/E311^{H}/E418^{-}$), and #5-1 ($E33^{H}/E311^{H}/E418^{-}$). RMSDs were calculated using THESEUS (1,2).

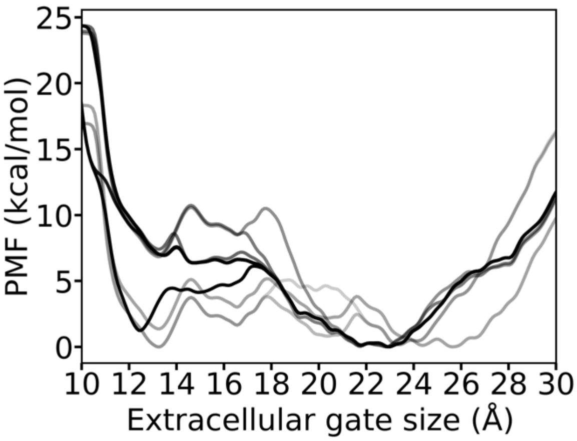


**Figure S6.** Time evolution of the potential of mean force of the extracellular gate size in a well-tempered metadynamics simulation. The transparency of the curve indicates the accumulative simulation time from 1 to 3 μs with a 200-ns spacing. The metadynamics biased both the extracellular and intracellular gate sizes, and used an initial Gaussian height = 0.6 kcal/mol. The bias factor was 25, Gaussian widths = (0.25 Å, 0.25 Å), and Gaussian were deposited every 100 ps.

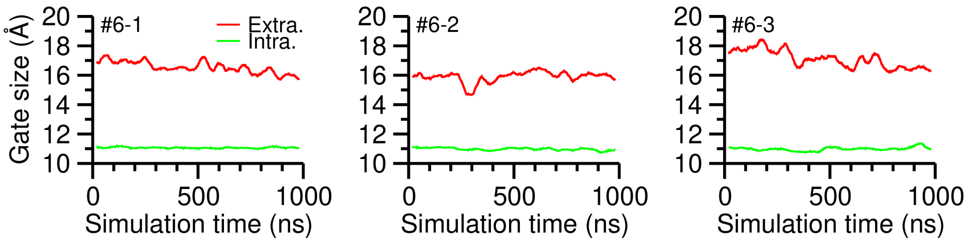


**Figure S7.** Time evolution of the extracellular (red lines) and intracellular (green lines) gate sizes in $E33^H/E311^-/E418^-$ (run #6) simulations initiated from the outward-facing state. A running average with a 20-ns window was performed on the time series.

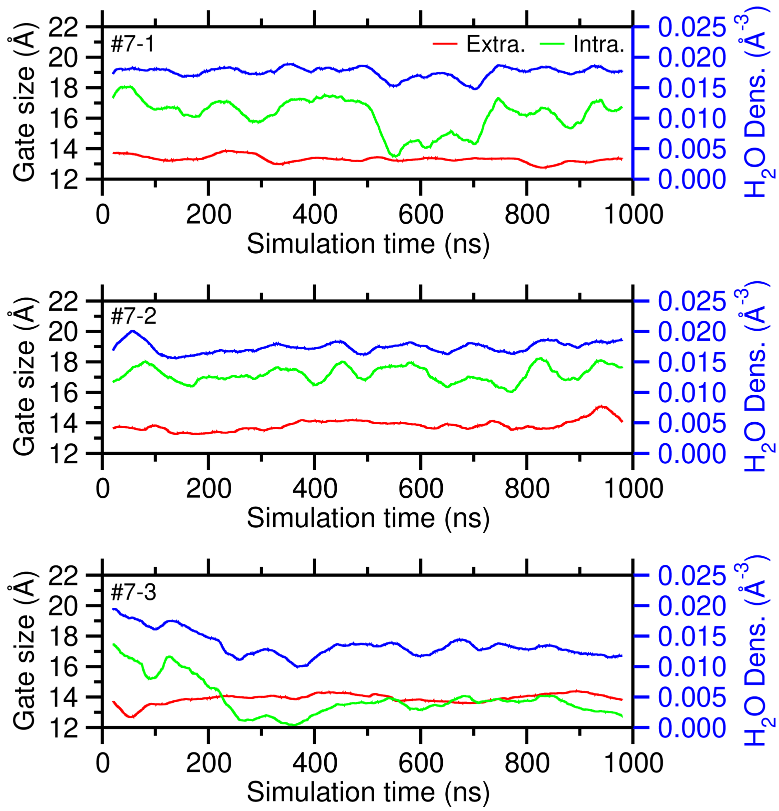


**Figure S8.** Time evolution of the extracellular (red lines) and intracellular (green lines) gate sizes and the water density around the intracellular gate in E33$^{-}$/E311$^{-}$/E418$^{-}$ (run #7) simulations. A running average using a 20-ns window was performed on the time series.

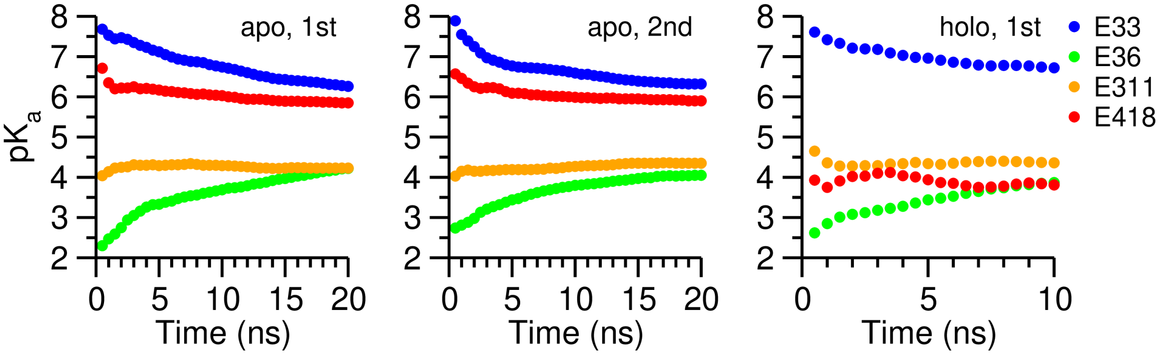


**Figure S9.** Convergence of the p$K_a$ values for E33, E36, E311, and E418 in $PepT_{Sh}$ calculated by pH-REX CpHMD. p$K_a$s were calculated cumulatively versus simulation time.

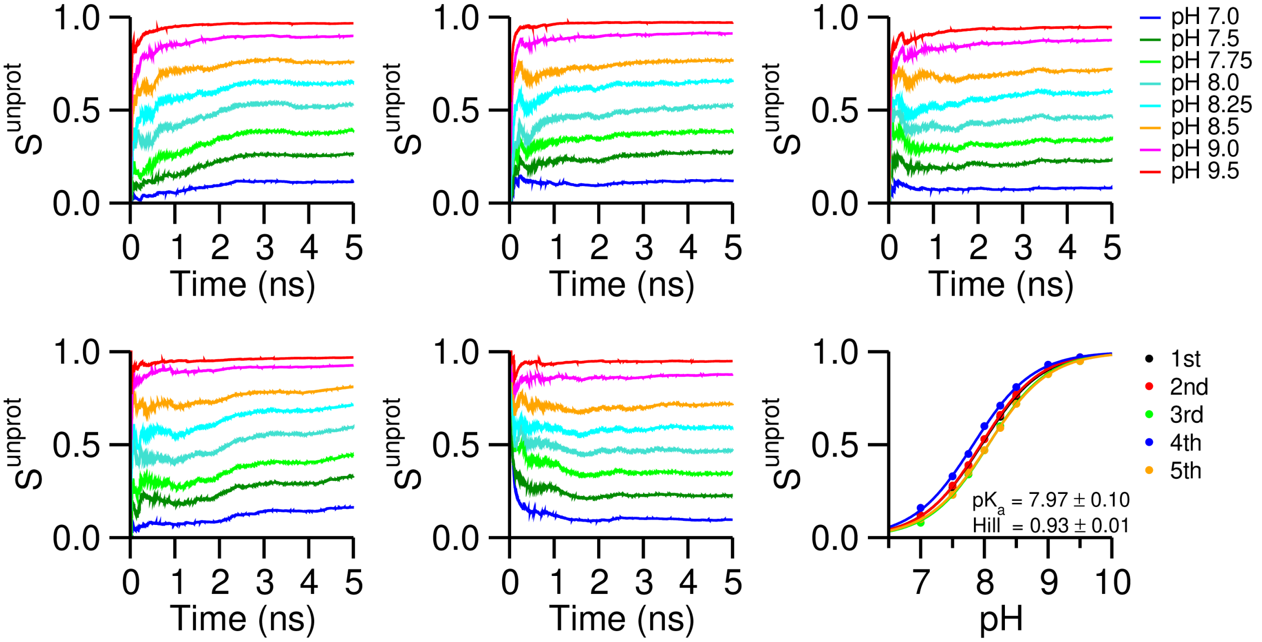


**Figure S10.** Validation of the hybrid-solvent CpHMD parameters for N-terminus (NT). To calculated the p$K_a$ of NT in aqueous, 5 independent pH-REX CpHMD simulations were performed. The titration curves are plotted in the bottom right panel with p$K_a$ and Hill coefficient $n$ reported (mean ± standard deviation). Other panels plot the convergence of the deprotonated fraction $S^{unprot}$ cumulatively calculated *vs.* time.

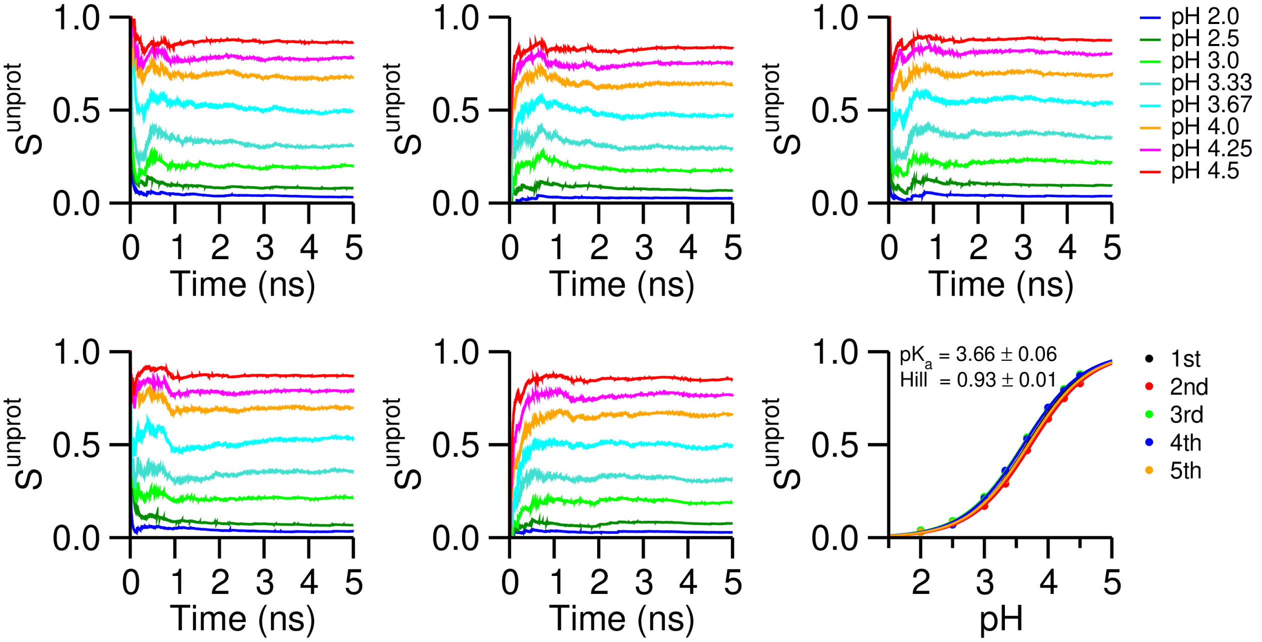


**Figure S11.** Validation of the hybrid-solvent CpHMD parameters for C-terminus (CT). To calculated the p$K_a$ of CT in aqueous, 5 independent pH-REX CpHMD simulations were performed. The titration curves are plotted in the bottom right panel with p$K_a$ and Hill coefficient $n$ reported (mean ± standard deviation). Other panels plot the convergence of the deprotonated fraction $S^{unprot}$ cumulatively calculated *vs.* time.

#### Table S1. Simulation details of classical MD

| Run ID | Protonation States (E33/E311/E418)[a] | Initial Configuration | Total Length (ns) | Data usage (time/Figure)[b] |
|---|---|---|---|---|
| 1-1 | +/-/+ | Crystal (6EXS) | 1000 | 0–1000 ns/Figure 2C<br>0–1000 ns/Figure 3B<br>0–1000 ns/ Figure S1<br>0–1000 ns/ Figure S2 |
| 1-2 | | | | 0–1000 ns/ Figure S1<br>0–1000 ns/ Figure S2 |
| 1-3 | | | | 0–1000 ns/ Figure S1<br>0–1000 ns/ Figure S2 |
| 2-1 | +/-/- | Crystal (6EXS) | 3500 | 0–1000 ns/Figure 2C<br>500 ns/Figure 3A<br>0–1000 ns/Figure 3B<br>0–1000 ns/Figure 4B<br>0–3500 ns/Figure 5ABC<br>2690.2 ns/Figure 5D<br>0–3500ns/Figure S1<br>0–3500ns/Figure S2<br>0–3500ns/Figure S3<br>0–3500ns/Figure S4<br>0-3500 ns/ Figure S5 |
| 2-2 | | | | 0–3500ns/Figure S1<br>0–3500ns/Figure S2<br>0–3500ns/Figure S3<br>0–3500ns/Figure S4 |
| 2-3 | | | | |

| | | | | |
|---|---|---|---|---|
| 3-1 | +/-/+ | 550 ns of #2-1 | 1000 | 0–1000 ns/Figure 4A<br>0–200 ns/Figure 4C |
| 3-2 | | 800 ns of #2-1 | | 0–1000 ns/Figure 4A<br>0–20 ns/Figure 4D |
| 3-3 | | 1050 ns of #2-1 | | |
| 4-1 | +/+/- | 2500 ns of #2-1 | 1000 | 100–500 ns/Figure 5AB<br>500 ns/Figure 5E<br>500 ns/Figure 6<br>0–1000 ns/ Figure S4<br>0–1000 ns/ Figure S5 |
| 4-2 | | 2450 ns of #2-1 | | 0–1000 ns/ Figure S4 |
| 4-3 | | 2500 ns of #2-1 | | |
| 5-1 | +/+/- | 3250 ns of #2-1 | 1000 | 0–500 ns/Figure 5AB<br>0–1000 ns/ Figure S4<br>0–1000 ns/ Figure S5 |
| 5-2 | | 2100 ns of #2-1 | | 0–1000 ns/ Figure S4 |
| 5-3 | | 3250 ns of #2-1 | | |
| 6-1 | +/-/- | 150 ns of #4-1 | 1000 | 0–1000 ns/ Figure S7 |
| 6-2 | | 250 ns of #4-1 | | |
| 6-3 | | 150 ns of #4-1 | | |
| 7-1 | -/-/- | Crystal (6EXS) | 1000 | 0–1000 ns/ Figure S8 |
| 7-2 | | | | |
| 7-3 | | | | |

[a] Protonation is indicated by "+", and deprotonation is indicated by "-". All the other ionizable residues were assigned their default protonation states except H22, H179, H187, H253, H399, and ligand termini, which were protonated according to CpHMD p$K_a$ calculations (Supplementary Table 3). [b] The time ranges or time points used for making specified figures.

#### Table S2. MS-RMD parameters for Glu

| | | | |
|---|---|---|---|
| $B$ | 1.94530 | $V_{ii}$ | -151.2996 |
| $b$ | 1.40003 | $\epsilon^{\mathrm{LJ}}_{\mathrm{OE-HH}}$ | 0.173646 |
| $b'$ | 1.08892 | $\sigma^{\mathrm{LJ}}_{\mathrm{OE-HH}}$ | 1.35219 |
| $C$ | 1.90167 | $\epsilon^{\mathrm{LJ}}_{\mathrm{Ow-HEP}}$ | 0.544690 |
| $c$ | 1.29037 | $\sigma^{\mathrm{LJ}}_{\mathrm{Ow-HEP}}$ | 1.35577 |
| $c_1$ | -25.0477 | $\epsilon^{\mathrm{LJ}}_{\mathrm{OE-OH}}$ | 0.160421 |
| $c_2$ | 2.95380 | $\sigma^{\mathrm{LJ}}_{\mathrm{OE-OH}}$ | 3.09726 |
| $c_3$ | 1.36184 | $\epsilon^{\mathrm{LJ}}_{\mathrm{OEP-Ow}}$ | 0.0775357 |
| $D$ | 143.003 | $\sigma^{\mathrm{LJ}}_{\mathrm{OEP-Ow}}$ | 3.06560 |
| $\alpha$ | 1.8 | | |
| $r_0$ | 0.975 | | |

The units of the listed parameters use kcal/mol as the energy unit and Å as the length unit.

**Table S3. p$K_a$s from pH-REX hybrid-solvent CpHMD**

| Residue | p$K_a$ values[a] | | |
|---|---|---|---|
| | Apo, 1st[b] | Apo, 2nd[c] | Holo[b] |
| D63 | 2.8 (0.9) | 2.5 (0.7) | 2.8 (0.9) |
| D90 | 3.9 (0.8) | 4.0 (0.9) | 4.0 (0.9) |
| D152 | 2.1 (0.9) | 2.1 (0.8) | 2.3 (1.0) |
| D156 | 3.6 (0.9) | 3.4 (1.0) | 3.5 (0.9) |
| D182 | 2.5 (0.8) | 2.2 (0.8) | 2.5 (0.9) |
| D258 | 2.5 (0.7) | 2.5 (0.9) | 2.4 (0.8) |
| D284 | 3.6 (0.9) | 3.7 (0.8) | 3.8 (0.9) |
| D287 | 3.5 (0.8) | 3.3 (0.8) | 2.7 (0.9) |
| D326 | 4.3 (0.8) | 4.4 (0.9) | 4.3 (0.8) |
| **E33** | **6.3 (0.6)** | **6.3 (0.7)** | **6.7 (0.6)** |
| **E36** | **4.2 (0.7)** | **4.0 (0.8)** | **3.9 (0.8)** |
| E150 | 3.7 (1.0) | 3.7 (1.0) | 3.8 (0.9) |
| E226 | 3.4 (0.9) | 3.3 (0.9) | 3.1 (0.9) |
| E227 | 2.3 (0.8) | 2.1 (0.9) | 2.4 (1.3) |
| E289 | 4.1 (0.8) | 4.2 (0.8) | 4.7 (0.9) |
| **E311** | **4.2 (1.0)** | **4.3 (1.0)** | **4.4 (0.8)** |
| E323 | 3.1 (0.8) | 3.1 (0.9) | 3.2 (0.9) |
| E340 | 4.7 (0.7) | 4.6 (0.8) | 4.7 (0.7) |
| **E418** | **5.8 (0.8)** | **5.9 (0.8)** | **3.8 (0.6)** |
| H22 | 7.7 (0.8) | 7.7 (0.8) | 7.4 (0.7) |
| H56 | 5.8 (0.8) | 5.7 (0.8) | 5.4 (0.8) |
| H110 | 5.2 (0.7) | 5.4 (0.7) | 5.3 (0.6) |
| H179 | 7.6 (0.9) | 7.6 (0.9) | 7.6 (0.9) |
| H187 | 7.8 (0.8) | 7.9 (0.8) | 7.8 (0.8) |
| H253 | 7.6 (0.8) | 7.7 (0.9) | 7.5 (0.8) |
| H399 | 7.1 (0.9) | 7.1 (0.9) | 7.1 (0.9) |
| K17 | 10.2 (0.9) | - | 10.3 (0.9) |
| K64 | Stay protonated | - | Stay protonated |
| **K137** | Stay protonated | - | Stay protonated |
| K210 | 10.4 (0.8) | - | 10.3 (0.8) |
| K225 | 10.3 (0.8) | - | 10.4 (0.8) |
| K228 | 9.9 (0.8) | - | 9.8 (0.7) |
| K230 | 11.4 (0.6) | - | 11.5 (0.5) |
| K283 | 10.7 (0.9) | - | 10.8 (0.9) |
| K294 | 9.4 (0.7) | - | 9.5 (0.8) |
| K364 | 10.7 (0.6) | - | 9.6 (0.5) |
| K367 | 10.2 (0.8) | - | 10.4 (0.8) |
| K368 | 10.3 (0.8) | - | 10.4 (0.8) |
| K375 | Stay protonated | - | Stay protonated |
| K431 | 10.1 (0.7) | - | 10.4 (0.8) |
| K435 | 10.6 (0.9) | - | 10.7 (0.9) |
| K461 | 11.3 (0.7) | - | 11.4 (0.6) |

| | | | |
|---|---|---|---|
| K464 | 10.5 (0.8) | - | 10.6 (0.8) |
| K492 | 10.0 (0.7) | - | 9.9 (0.7) |
| K495 | 10.4 (0.8) | - | 10.3 (0.7) |
| R24 | Stay protonated | - | Stay protonated |
| **R37** | Stay protonated | - | Stay protonated |
| R44 | Stay protonated | - | Stay protonated |
| R91 | Stay protonated | - | Stay protonated |
| R96 | Stay protonated | - | Stay protonated |
| R146 | Stay protonated | - | Stay protonated |
| R154 | Stay protonated | - | Stay protonated |
| R184 | Stay protonated | - | Stay protonated |
| R209 | Stay protonated | - | Stay protonated |
| R229 | Stay protonated | - | Stay protonated |
| R281 | Stay protonated | - | Stay protonated |
| R290 | Stay protonated | - | Stay protonated |
| R292 | Stay protonated | - | Stay protonated |
| R324 | Stay protonated | - | Stay protonated |
| R337 | Stay protonated | - | Stay protonated |
| C112 | 11.1 (0.6) | - | 11.1 (0.6) |
| C414 | 10.9 (0.8) | - | 11.6 (1.3) |
| C420 | 9.5 (0.5) | - | Stay protonated |
| Y40 | Stay protonated | - | Stay protonated |
| Y41 | 11.3 (0.6) | - | Stay protonated |
| Y50 | Stay protonated | - | Stay protonated |
| Y52 | 11.0 (0.7) | - | 11.1 (0.8) |
| Y74 | Stay protonated | - | Stay protonated |
| Y79 | 10.5 (0.7) | - | 11.3 (0.9) |
| Y148 | Stay protonated | - | Stay protonated |
| Y163 | 10.7 (0.8) | - | 10.4 (1.0) |
| Y204 | Stay protonated | - | Stay protonated |
| Y231 | 11.4 (0.4) | - | 11.1 (0.3) |
| Y250 | Stay protonated | - | Stay protonated |
| Y251 | Stay protonated | - | Stay protonated |
| Y275 | 11.5 (0.6) | - | 11.5 (0.9) |
| Y320 | Stay protonated | - | Stay protonated |
| Y387 | Stay protonated | - | Stay protonated |
| Y397 | 11.2 (0.6) | - | 10.9 (0.7) |
| Y411 | Stay protonated | - | Stay protonated |
| Y471 | Stay protonated | - | Stay protonated |
| NT-CSM[d] | 16.0 (0.9) | | Stay protonated |
| CT-GLY[d] | 8.6 (0.9) | | 9.8 (0.6) |

[a] Parenthesized are Hill coefficients. “Stay protonated” indicates a residue, though ionizable in CpHMD, did not titrate within the pH range studied. [b] pH range 2.0–11.75. [c] pH range 2.0–9.0. Lys/Arg/Cys/Tyr permanently protonated. [d] pH range 7.0–16.5.

#### Supporting References


1. Theobald, D. L., and D. S. Wuttke. 2008. Accurate Structural Correlations from Maximum Likelihood Superpositions. *PLoS Comput. Biol.* 4:e43.
2. Theobald, D. L., and P. A. Steindel. 2012. Optimal simultaneous superpositioning of multiple structures with missing data. *Bioinformatics* 28:1972−1979.

### Appendix I: CHARMM topology and parameter for S-Cys-3M3SH

```
* Toppar stream file generated by
* CHARMM General Force Field (CGenFF) program version 2.2.0
* For use with CGenFF version 4.0
* To model dipeptide S-Cys-Gly-3M3SH, define S-Cys-3M3SH a new non-standard amino acid “CSM”.
* Check Minhas et al., eLife, 2018, 7: e34995
* Generated by Zhi (Shane) Yue, Gregory A. Voth lab, UChicago, 02/18/2019
*

read rtf card append
* Topologies generated by
* CHARMM General Force Field (CGenFF) program version 2.2.0
*
36 1

! "penalty" is the highest penalty score of the associated parameters.
! Penalties lower than 10 indicate the analogy is fair; penalties between 10
! and 50 mean some basic validation is recommended; penalties higher than
! 50 indicate poor analogy and mandate extensive validation/optimization.

DECL -CA
DECL -C
DECL -O
DECL +N
DECL +HN
DECL +CA

RESI CSM            0.000
GROUP
ATOM N      NH1    -0.470 ! from C22 CYS
ATOM HN     H       0.310 ! from C22 CYS
ATOM CA     CT1     0.070 ! from C22 CYS
ATOM HA     HB      0.090 ! from C22 CYS
GROUP
ATOM CB     CT2    -0.132 ! from C22 CYS, ATOM CHARGE adjusted for charge neutrality
ATOM HB1    HA      0.090 ! from C22 CYS
ATOM HB2    HA      0.090 ! from C22 CYS
ATOM SG     S      -0.092 ! penalty = 21.575 from MSH CGENFF, ATOM TYPE replaced by C22 CYS
ATOM CD     CG301   0.238 ! penalty = 27.831 from MSH CGENFF
ATOM CE1    CG331  -0.331 ! penalty = 10.272 from MSH CGENFF
ATOM HE11   HGA3    0.090 ! penalty =  0.810 from MSH CGENFF
ATOM HE12   HGA3    0.090 ! penalty =  0.810 from MSH CGENFF                 HE12
ATOM HE13   HGA3    0.090 ! penalty =  0.810 from MSH CGENFF                 |
ATOM CE2    CG321  -0.246 ! penalty = 14.778 from MSH CGENFF     |      HE11-CE1-HE13
ATOM HE21   HGA2    0.090 ! penalty =  0.900 from MSH CGENFF  HN-N           |
ATOM HE22   HGA2    0.090 ! penalty =  0.900 from MSH CGENFF     |   HB1     |   HE21  HZ21  HH21
ATOM CZ2    CG321  -0.180 ! penalty =  0.901 from MSH CGENFF     |   |       |   |     |     |
ATOM HZ21   HGA2    0.090 ! penalty =  0.000 from MSH CGENFF  HA-CA--CB--SG--CD--CE2---CZ2---CH2-HH22
ATOM HZ22   HGA2    0.090 ! penalty =  0.000 from MSH CGENFF     |   |       |   |     |     |
ATOM CH2    CG331  -0.270 ! penalty =  0.045 from MSH CGENFF     |   HB2     |   HE22  HZ22  HH23
ATOM HH21   HGA3    0.090 ! penalty =  0.000 from MSH CGENFF   O=C           |
ATOM HH22   HGA3    0.090 ! penalty =  0.000 from MSH CGENFF     |      HE31-CE3-HE32
ATOM HH23   HGA3    0.090 ! penalty =  0.000 from MSH CGENFF                 |
ATOM CE3    CG321  -0.249 ! penalty = 14.786 from MSH CGENFF            HZ31-CZ3-HZ32
ATOM HE31   HGA2    0.090 ! penalty =  0.900 from MSH CGENFF                 |
ATOM HE32   HGA2    0.090 ! penalty =  0.900 from MSH CGENFF                 OH3
ATOM CZ3    CG321   0.051 ! penalty =  1.065 from MSH CGENFF                  \
ATOM HZ31   HGA2    0.090 ! penalty =  0.000 from MSH CGENFF                   HH3
ATOM HZ32   HGA2    0.090 ! penalty =  0.000 from MSH CGENFF
ATOM OH3    OG311  -0.649 ! penalty =  0.090 from MSH CGENFF
ATOM HH3    HGP1    0.420 ! penalty =  0.000 from MSH CGENFF
GROUP
ATOM C      C       0.510 ! from C22 CYS
ATOM O      O      -0.510 ! from C22 CYS

BOND N   HN
BOND N   CA
BOND CA  HA
BOND C   CA
BOND C   +N
BOND CA  CB
BOND CB  HB1
BOND CB  HB2
BOND CB  SG
BOND SG  CD
BOND CD  CE1
BOND CE1 HE11
BOND CE1 HE12
BOND CE1 HE13
BOND CD  CE2
BOND CE2 HE21
BOND CE2 HE22
BOND CE2 CZ2
BOND CZ2 HZ21
BOND CZ2 HZ22
BOND CZ2 CH2
BOND CH2 HH21
BOND CH2 HH22
BOND CH2 HH23
BOND CD  CE3
BOND CE3 HE31
BOND CE3 HE32
BOND CE3 CZ3
BOND CZ3 HZ31
BOND CZ3 HZ32
BOND CZ3 OH3
BOND OH3 HH3
DOUBLE C O
IMPR N -C CA HN  C CA +N O
CMAP -C  N  CA  C   N  CA  C  +N
DONOR HN N
DONOR HH3 OH3
ACCEPTOR O C
IC -C   CA   *N   HN    1.3479 123.9300  180.0000 114.7700  0.9982 ! from C22 CYS
IC -C   N    CA   C     1.3479 123.9300  180.0000 105.8900  1.5202 ! from C22 CYS
IC N    CA   C    +N    1.4533 105.8900  180.0000 118.3000  1.3498 ! from C22 CYS
IC +N   CA   *C   O     1.3498 118.3000  180.0000 120.5900  1.2306 ! from C22 CYS
IC CA   C    +N   +CA   1.5202 118.3000  180.0000 124.5000  1.4548 ! from C22 CYS
IC N    C    *CA  CB    1.4533 105.8900  121.7900 111.9800  1.5584 ! from C22 CYS
IC N    C    *CA  HA    1.4533 105.8900 -116.3400 107.7100  1.0837 ! from C22 CYS
IC N    CA   CB   SG    1.4533 111.5600  180.0000 113.8700  1.8359 ! from C22 CYS
IC SG   CA   *CB  HB1   1.8359 113.8700  119.9100 107.2400  1.1134 ! from C22 CYS
IC SG   CA   *CB  HB2   1.8359 113.8700 -125.3200 109.8200  1.1124 ! from C22 CYS
IC CA   CB   SG   CD    1.5460 110.2800  180.0000  98.9400  1.8180 ! from C22 MET CB-CG-SD-CE, S-CG301 updated
```

```
IC CB   SG   CD   CE1   1.8180  95.0000  180.0000 114.5000  1.5380 ! CT2-S,       CT2-S-CG301,       CT2-S-CG301-CG331,       S-CG301-CG331,     CG301-CG331
IC CE1  SG   *CD  CE2   1.5380 114.5000  120.0000 114.5000  1.5380 ! CG331-CG301, CG331-CG301-S,     CG331-S-CG301-CG321,     S-CG301-CG321,     CG301-CG321
IC CE1  SG   *CD  CE3   1.5380 114.5000 -120.0000 114.5000  1.5380 ! CG331-CG301, CG331-CG301-S,     CG331-S-CG301-CG321,     S-CG301-CG321,     CG301-CG321
IC SG   CD   CE1  HE11  1.8180 114.5000    0.0000 110.1000  1.1110 ! S-CG301,     S-CG301-CG331,     S-CG301-CG331-HGA3,      CG301-CG331-HGA3,  CG331-HGA3
IC HE11 CD   *CE1 HE12  1.1110 110.1000  120.0000 110.1000  1.1110 ! HGA3-CG331,  HGA3-CG331-CG301,  HGA3-CG301-CG331-HGA3,   CG301-CG331-HGA3,  CG331-HGA3
IC HE11 CD   *CE1 HE13  1.1110 110.1000 -120.0000 110.1000  1.1110 ! HGA3-CG331,  HGA3-CG331-CG301,  HGA3-CG301-CG331-HGA3,   CG301-CG331-HGA3,  CG331-HGA3
IC SG   CD   CE2  CZ2   1.8180 114.5000    0.0000 113.5000  1.5300 ! S-CG301,     S-CG301-CG321,     S-CG301-CG321-CG321,     CG301-CG321-CG321, CG321-CG321
IC CZ2  CD   *CE2 HE21  1.5300 113.5000  120.0000 110.1000  1.1110 ! CG321-CG321, CG321-CG321-CG301, CG321-CG301-CG321-HGA2,  CG301-CG321-HGA2,  CG321-HGA2
IC CZ2  CD   *CE2 HE22  1.5300 113.5000 -120.0000 110.1000  1.1110 ! CG321-CG321, CG321-CG321-CG301, CG321-CG301-CG321-HGA2,  CG301-CG321-HGA2,  CG321-HGA2
IC CD   CE2  CZ2  CH2   1.5380 113.5000    0.0000 115.0000  1.5280 ! CG301-CG321, CG301-CG321-CG321, CG301-CG321-CG321-CG331, CG321-CG321-CG331, CG321-CG331
IC CH2  CE2  *CZ2 HZ21  1.5280 115.0000  120.0000 110.1000  1.1110 ! CG331-CG321, CG331-CG321-CG321, CG331-CG321-CG321-HGA2,  CG321-CG321-HGA2,  CG321-HGA2
IC CH2  CE2  *CZ2 HZ22  1.5280 115.0000 -120.0000 110.1000  1.1110 ! CG331-CG321, CG331-CG321-CG321, CG331-CG321-CG321-HGA2,  CG321-CG321-HGA2,  CG321-HGA2
IC CE2  CZ2  CH2  HH21  1.5300 115.0000    0.0000 110.1000  1.1110 ! CG321-CG321, CG321-CG321-CG331, CG321-CG321-CG331-HGA3,  CG321-CG331-HGA3,  CG331-HGA3
IC HH21 CZ2  *CH2 HH22  1.1110 110.1000  120.0000 110.1000  1.1110 ! HGA3-CG331,  HGA3-CG331-CG321,  HGA3-CG321-CG331-HGA3,   CG321-CG331-HGA3,  CG331-HGA3
IC HH21 CZ2  *CH2 HH23  1.1110 110.1000 -120.0000 110.1000  1.1110 ! HGA3-CG331,  HGA3-CG331-CG321,  HGA3-CG321-CG331-HGA3,   CG321-CG331-HGA3,  CG331-HGA3
IC SG   CD   CE3  CZ3   1.8180 114.5000    0.0000 113.5000  1.5300 ! S-CG301,     S-CG301-CG321,     S-CG301-CG321-CG321,     CG301-CG321-CG321, CG321-CG321
IC CZ3  CD   *CE3 HE31  1.5300 113.5000  120.0000 110.1000  1.1110 ! CG321-CG321, CG321-CG321-CG301, CG321-CG301-CG321-HGA2,  CG301-CG321-HGA2,  CG321-HGA2
IC CZ3  CD   *CE3 HE32  1.5300 113.5000 -120.0000 110.1000  1.1110 ! CG321-CG321, CG321-CG321-CG301, CG321-CG301-CG321-HGA2,  CG301-CG321-HGA2,  CG321-HGA2
IC CD   CE3  CZ3  OH3   1.5380 113.5000    0.0000 110.1000  1.4200 ! CG301-CG321, CG301-CG321-CG321, CG301-CG321-CG321-OG311, CG321-CG321-OG311, CG321-OG311
IC OH3  CE3  *CZ3 HZ31  1.4200 110.1000  120.0000 110.1000  1.1110 ! OG311-CG321, OG311-CG321-CG321, OG311-CG321-CG321-HGA2,  CG321-CG321-HGA2,  CG321-HGA2
IC OH3  CE3  *CZ3 HZ32  1.4200 110.1000 -120.0000 110.1000  1.1110 ! OG311-CG321, OG311-CG321-CG321, OG311-CG321-CG321-HGA2,  CG321-CG321-HGA2,  CG321-HGA2
IC CE3  CZ3  OH3  HH3   1.5300 110.1000    0.0000 106.0000  0.9600 ! CG321-CG321, CG321-CG321-OG311, CG321-CG321-OG311-HGP1,  CG321-OG311-HGP1,  OG311-HGP1

END

read param card flex append
* Parameters generated by analogy by
* CHARMM General Force Field (CGenFF) program version 2.2.0
*

! Penalties lower than 10 indicate the analogy is fair; penalties between 10
! and 50 mean some basic validation is recommended; penalties higher than
! 50 indicate poor analogy and mandate extensive validation/optimization.

BONDS
CG301  S       198.00     1.8180 ! C3H, from CG321 SG311, penalty = 12, SG311 replaced by S

ANGLES
CG321  CG301  S        58.00    114.50 ! C3H, from CG321 CG321 SG311, penalty = 12,  SG311 replaced by S
CG331  CG301  S        58.00    114.50 ! C3H, from CG331 CG321 SG311, penalty = 12,  SG311 replaced by S
CG301  S      CT2      34.00     95.00 ! C3H, from CG321 SG311 CG321, penalty = 1.8, SG311 replaced by S, CG321 replaced by CT2

DIHEDRALS
S      CG301  CG321  CG321    0.1950  3   0.00 ! C3H, from CG321 CG321 CG321 SG311,  penalty = 12,   SG311 replaced by S
S      CG301  CG321  HGA2     0.0100  3   0.00 ! C3H, from SG311 CG321 CG321 HGA2,   penalty = 12,   SG311 replaced by S
S      CG301  CG331  HGA3     0.1600  3   0.00 ! C3H, from SG311 CG321 CG331 HGA3,   penalty = 12,   SG311 replaced by S
CG321  CG301  S      CT2      0.2400  1 180.00 ! C3H, from CG321 CG321 SG311 CG321,  penalty = 12,   SG311 replaced by S, CG321 replaced by CT2
CG321  CG301  S      CT2      0.3700  3   0.00 ! C3H, from CG321 CG321 SG311 CG321,  penalty = 12,   SG311 replaced by S, CG321 replaced by CT2
CG331  CG301  S      CT2      0.2400  1 180.00 ! C3H, from CG321 CG321 SG311 CG321,  penalty = 12.9, SG311 replaced by S, CG321 replaced by CT2
CG331  CG301  S      CT2      0.3700  3   0.00 ! C3H, from CG321 CG321 SG311 CG321,  penalty = 12.9, SG311 replaced by S, CG321 replaced by CT2
CG301  CG321  CG321  CG331    0.1950  3   0.00 ! C3H, from CG301 CG321 CG321 CG321,  penalty = 0.9
CG301  CG321  CG321  OG311    0.1950  3   0.00 ! C3H, from CG321 CG321 CG321 OG311,  penalty = 1.8
CT1    CT2    S      CG301    0.1950  3   0.00 ! C3H, from CG324 CG321 SG311 CG321,  penalty = 2.4,  SG311 replaced by S, CG321 replaced by CT2, CG314 replaced by CT1
HA     CT2    S      CG301    0.2800  3   0.00 ! C3H, from HGA2  CG321 SG311 CG321,  penalty = 1.8,  SG311 replaced by S, CG321 replaced by CT2, HGA2  replaced by HA

IMPROPERS

END
RETURN
```

#### Appendix II: CHARMM hybrid-solvent CpHMD parameters for N-terminus

```
* Parameter file for HPHMD
* Charges generated based on C22 PRES NTER & NNEU
* Parameterized by Zhi (Shane) Yue, Gregory A. Voth lab, UChicago, February 2019
* Syntax:
* RESNAME EXPT_PKA PARA PARB BARR
*  ATOM_NAME PROT_CHARGE UNPROT_CHARGE PROT_RAD UNPROT_RAD
*

! -------------------------
! Experimental pKa: 8.00 +/- 0.03
! Measured at 298.15 K, 1 atm, 0.1 M NaCl using pentapeptide NH3+-AAAAA-CONH2
! Check Thurlkill et al., Protein Sci.,2006, 15(5): 1214-1218
! -------------------------
! 5 independent pH-REX hybrid-solvent CpHMD runs
!
! CHARMM version: c42b2
! C22+CMAP CHARMM Force Field
! Truncated octahedron box, 39 Angstrom
! 1 NTER-AAAAA-CT2, 1463 CHARMM-modified TIP3P waters, no ions
! 300.00 K (Nosé-Hoover), 1 atm (Langevin piston)
!
! GB settings:
! gbsw hybrid sgamma 0.000 nang 50 conc 0.000 temp 300.00 -
!      sele SOLUTE end
!
! Nonbonded settings:
! nbond elec atom cdie vdw vatom vswitch -
!       ctonnb 10.0 ctofnb 12.0 cutnb 14.0 cutim 16.0 -
!       ewald pmew fftx 40 ffty 40 fftz 40 kappa 0.34 spline order 6 -
!       inbfrq -1 imgfrq -1
! -------------------------
! pKa(calc) = 7.97 +/- 0.10
! n  (calc) = 0.93 +/- 0.01
!
! Fraction of mixed state: <= 24%
! -------------------------
! To use with other terminal residues such as NH3+-XXX-,
! make a copy of the entry below and replace “Ala” with “XXX”
! -------------------------

NTALA 8.00 -78.5636 1.04627435067512 1.5
N    -0.30  -0.97         !     HT1                     HT1
HT1   0.33   0.33         !    /                       /
HT2   0.33   0.33         ! --N--HT3* (+)  <=====>  --N
HT3   0.33   0.00 1.0 0.0 !    \                       \
                          !     HT2                     HT2
END
```

#### Appendix III: CHARMM hybrid-solvent CpHMD parameters for C-terminus

```
* Parameter file for HPHMD
* Charges generated based on C22 PRES CTER & CNEU
* Parameterized by Zhi (Shane) Yue, Gregory A. Voth lab, UChicago, February 2019
* Syntax:
* RESNAME EXPT_PKA PARA PARB BARR
*  ATOM_NAME PROT_CHARGE UNPROT_CHARGE PROT_RAD UNPROT_RAD
* RESNAME EXPT_PKA PARA PARB BARR PARA10 PARB10 BARTAU
*                  R1   R2   R3   R4     R5     R6
*  ATOM_NAME PROT_CHARGE UNPROT_CHARGE PROT_RAD UNPROT_RAD
*

! -------------------------
! Experimental pKa: 3.67 +/- 0.03
! Measured at 298.15 K, 1 atm, 0.1 M NaCl using pentapeptide CH3CONH-AAAAA-COO-
! Check Thurlkill et al., Protein Sci.,2006, 15(5): 1214-1218
! -------------------------
! 5 independent pH-REX hybrid-solvent CpHMD runs
!
! CHARMM version: c42b2
! C22+CMAP CHARMM Force Field
! Truncated octahedron box, 41 Angstrom
! 1 ACE-AAAAA-CTRP2, 1772 CHARMM-modified TIP3P waters, no ions
! 300.00 K (Nosé-Hoover), 1 atm (Langevin piston)
!
! GB settings:
! gbsw hybrid sgamma 0.000 nang 50 conc 0.000 temp 300.00 -
!      sele SOLUTE end
!
! Nonbonded settings:
! nbond elec atom cdie vdw vatom vswitch -
!       ctonnb 10.0 ctofnb 12.0 cutnb 14.0 cutim 16.0 -
!       ewald pmew fftx 40 ffty 40 fftz 40 kappa 0.34 spline order 6 -
!       inbfrq -1 imgfrq -1
! -------------------------
! pKa(calc) = 3.66 +/- 0.06
! n  (calc) = 0.93 +/- 0.01
!
! Fraction of mixed state: <= 30%
! -------------------------
! To make C-terminus titratable in CpHMD, apply patch “PRES CTRP2”
! Check documentation phmd.doc for CpHMD topology & parameter
! -------------------------
! To use with other terminal residues such as -XXX-COO-,
! make a copy of the entry below and replace “Ala” with “XXX”
! -------------------------

CTALA1 3.67 -90.202 0.183009 2.0
C      0.72   0.34         !    OT1-HC1*              OT1 (-)
OT1   -0.61  -0.67         !   /                     /
OT2   -0.55  -0.67         ! --C          <=====>  --C
HC1    0.44   0.00 1.0 0.0 !   \\                    \\
HC2    0.00   0.00         !    OT2                   OT2

CTALA2 3.67 -90.2589 0.182468 2.0 -29.8537 0.498661
            -26.7031 56.9114 -29.9575 0.490501 -90.1841 0.180540580420495
C      0.72   0.34         !    OT1                   OT1
OT1   -0.55  -0.67         !   //                    //
OT2   -0.61  -0.67         ! --C          <=====>  --C
HC1    0.00   0.00         !   \                     \
HC2    0.44   0.00 1.0 0.0 !    OT2-HC2*              OT2 (-)

END
```
